# Distinct H3K9me3 heterochromatin maintenance dynamics govern different gene programs and repeats in pluripotent cells

**DOI:** 10.1101/2024.09.16.613328

**Authors:** Jingchao Zhang, Greg Donahue, Michael B. Gilbert, Tomer Lapidot, Dario Nicetto, Kenneth S. Zaret

## Abstract

H3K9me3-heterochromatin, established by lysine methyltransferases (KMTs) and compacted by HP1 isoforms, represses alternative lineage genes and DNA repeats. Our understanding of H3K9me3-heterochromatin stability is presently limited to individual domains and DNA repeats. We engineered *Suv39h2* KO mouse embryonic stem cells to degrade remaining two H3K9me3- KMTs within one hour and found that both passive dilution and active removal contribute to H3K9me3 decay within 12-24 hours. We discovered four different H3K9me3 decay rates across the genome and chromatin features and transcription factor binding patterns that predict the stability classes. A “binary switch” governs heterochromatin compaction, with HP1 rapidly dissociating from heterochromatin upon KMTs’ depletion and a particular threshold level of HP1 limiting pioneer factor binding, chromatin opening, and exit from pluripotency within 12 hr. Unexpectedly, receding H3K9me3 domains unearth residual HP1β peaks enriched with heterochromatin-inducing proteins. Our findings reveal distinct H3K9me3-heterochromatin maintenance dynamics governing gene networks and repeats that together safeguard pluripotency.

## Introduction

Cells maintain their identity by physically compacting alternative lineage genes into various forms of transcriptionally silent heterochromatin^1^. Of the major forms of heterochromatin in pluripotent cells, heterochromatin marked by histone H3 lysine 9 trimethylation (H3K9me3) is necessary to maintain pluripotency and for cell differentiation^2–4^, while heterochromatin marked by H3K27me3 or DNA methylation is needed for differentiation but not for mouse embryonic stem cell (ESC) self-renewal^5,6^. H3K9me3 is bound by Heterochromatin protein 1 (HP1) isoforms to compact genomic domains into heterochromatin, restricting access by transcription factors and RNA polymerase to silence lineage-inappropriate genes and DNA repeats^7^. During DNA replication, parental histones in heterochromatin are recycled^8^ and H3K9me3 marks are recognized by chromodomains in “writer” KMTs SUV39H1 and SUV39H2, and the Tudor domain in the third “writer”, SETDB1^9^, and “reader” proteins HP1α/β/γ, which further recruit writers to modify newly loaded histones^10,11^. The reader-writer self-enforcing mechanism restores H3K9me3 heterochromatin in daughter cells, providing epigenetic inheritance that maintains cell identity^12^. However, the HP1 chromodomain has low affinity to the H3K9me3 mark in vitro and HP1 binding to heterochromatin is highly dynamic in vivo^13–15^. Proteomic quantifications of parental and naïve histones also reveal a slow restoration of H3K9me3 and H3K27me3 heterochromatin marks after DNA replication^16^. These points raise questions about the basis for H3K9me3 heterochromatin stability, how H3K9me3 maintenance may differ across the genome, and features that could impart differential stabilities.

Our understanding of heterochromatin maintenance has been dominated by single-locus studies, in which H3K9me3-heterochromatin is initiated by recruiting H3K9me3 machineries to a single ectopic locus^17–19^, leading to the spreading of heterochromatin and repression of nearby genes. In fission yeast, such an ectopic H3K9me2 domain is slowly eroded by the H3K9 demethylase (KDM) ortholog, Epe1, in a cell-cycle independent manner, leading to de-repression of a reporter gene within 10 cell generations^17,18^. Thus, yeast maintains ectopic heterochromatin domains with a balance between H3K9me3 KMTs and KDMs. Whether the mechanisms revealed by a single locus can be extrapolated to mammalian genomes remains to be assessed. Furthermore, ectopic heterochromatin experiments and the subsequent mathematic models assume a neutral state across chromatin, without accounting for different local chromatin environments that may modulate H3K9me3-heterochromatin maintenance^20–24^. Therefore, studying genome-wide H3K9me3 maintenance in mammalian cells necessitates global perturbations of endogenous H3K9me3-heterochromatin and assessing the consequences on diverse local chromatin states.

Recent studies have completely depleted H3K9me3 in chromatin by genetically ablating all H3K9me3 methyltransferases (KMTs) in mouse fibroblast or liver lineages, thereby activating alternative lineage genes and transposable elements^4,25,26^. However, the slow process for complete genetic deletion makes it difficult to distinguish direct versus indirect consequences. Using a combination of degron technologies^27,28^, we depleted all three H3K9me3 KMTs in pluripotent cells within one hour. Leveraging the acute degradation system, we assessed the immediate impact of KMT and H3K9me3 loss on the effector protein HP1, other heterochromatin-associated proteins, chromatin compaction, and the expression of different gene networks and repeat families at high temporal resolution. Our results reveal distinct H3K9me3-heterochromatin maintenance types for different gene networks and repeats in mammalian cells that together maintain pluripotency, and greatly expand our understanding of principles of H3K9me3 maintenance and remodeling beyond that discerned from single locus studies or conventional genetic deletions^17–19^.

## Results

### Acute degradation of KMTs reveals dynamic H3K9me3 maintenance

To investigate H3K9me3-heterochromatin dynamics, we developed a triple conditional knockout (cTKO) mouse ESC line from relevant mouse embryos^4^ (**Supplementary Fig. 1a-c**) and sequentially and homozygously tagged *Suv39h1* and *Setdb1* with IAA7 (auxin inducible degradation)^27^ and dTAG^28^, respectively, in a *Suv39h2* homozygous null background, to create “dTKO” cells, for **d**egradable **T**riple **K**nock-**O**ut. (**Fig. 1a and Supplementary Fig. 2a-f**). Sequences for the adaptor protein atAFB2, for auxin-inducible degradation^27^, were knocked into *TIGRE* “safe harbor” locus^29^. The dTKO mouse ESCs express pluripotent markers Sox2 and Esrrb in ES self-renewal condition (**Extended Data Fig. 1a**) and have been cultured for more than 40 passages. Furthermore, dTKO cells activate early lineage markers during embryoid body differentiation (**Extended Data Fig. 1b-c**), indicating that they remain pluripotent. Importantly, the H3K9me3 ChIP signals and the expression of various transposable elements in the dTKO cells correlate well with the parental cTKO cells (**Extended Data. Fig. 1d-e**), further validating that protein tagging does not affect the KMTs’ activities.

**Fig. 1:**
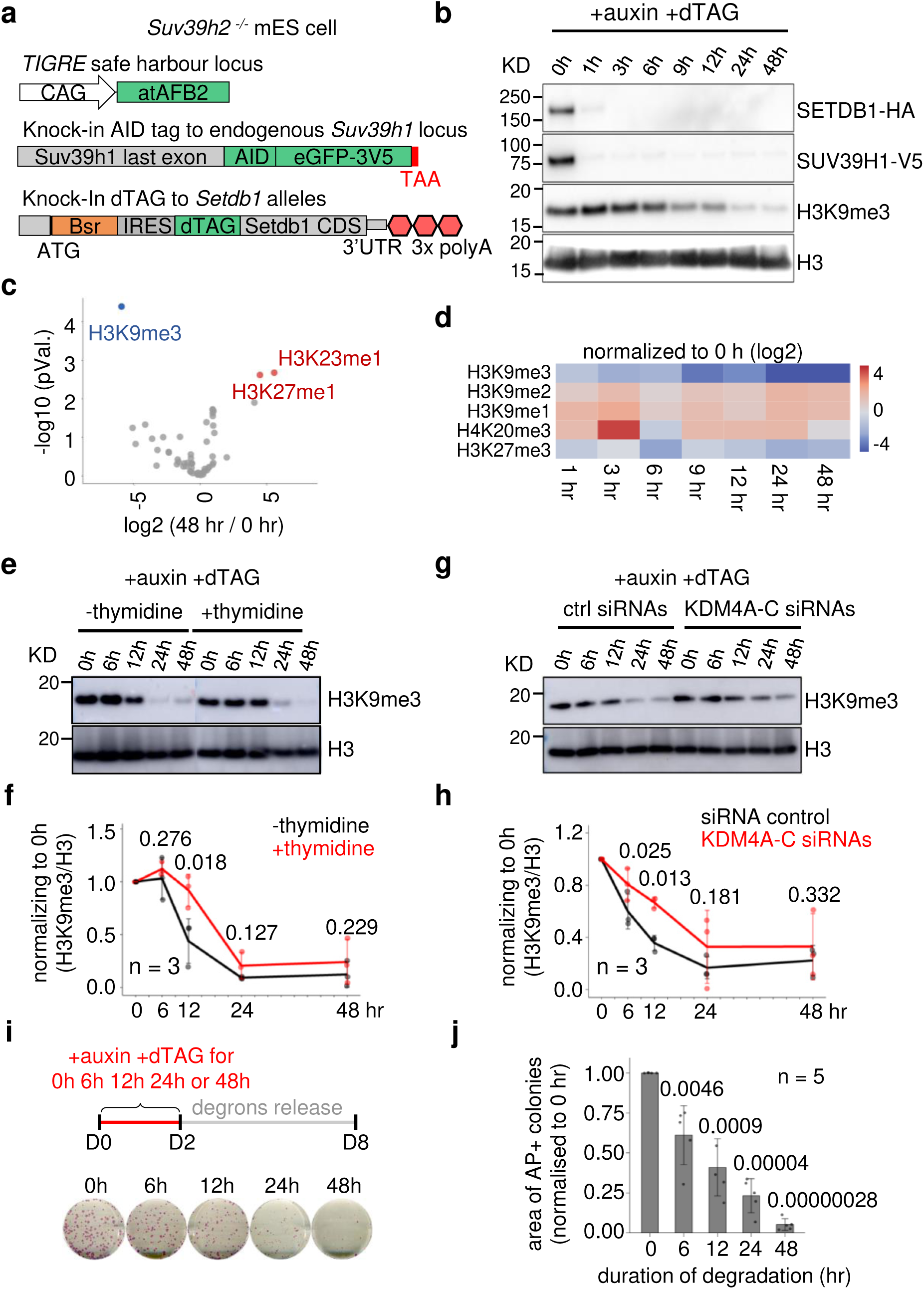
Acute depletion of H3K9me3 KMTs causes rapid decay of H3K9me3 marks in mouse ESCs **a**, Schematics of the genetic engineering of dTKO mouse ESC line. **b**, Western blot of SETDB1- HA and SUV39H1-V5 with designated tag antibodies, H3K9me3, and histone H3 in chromatin fraction after different hours of auxin and dTAG13 degrons treatment. Experiments were repeated three times independently with similar results. The molecular weight in KD (kilodaltons) is on the left. **c**, Differential analysis of the histone marks abundances measured by mass spectrometry between 0 hr and 48 hr of auxin and dTAG degrons treatment. Statistically significantly changed histone marks were highlighted in blue (reduced) and red (increased). **d**, Heatmap showing the dynamics of heterochromatin marks over the degradation time course. Values are log2 fold changes of the mean values of three biological replicates at each time point over 0 hr. **e,f,** Western blot (**e**) and quantifications (**f**) of H3K9me3 normalized to 0 hr sample after histone H3 normalization, following the auxin and dTAG treatments with or without thymidine; *n* = 3 biologically independent replicates. **g,h,** Western blot (**g**) and quantifications (**h**) of H3K9me3, following the auxin and dTAG treatment with non-targeting siRNAs (ctrl) or KDM4A-C siRNAs. H3K9me3 levels are normalized to 0 hr sample after histone H3 normalization; *n* = 3 biologically independent replicates. **i**, Schematics of different regimes of auxin and dTAG treatment of moue dTKO ESCs followed by degrons washout, before alkaline phosphatases staining (left). Quantifications of areas of alkaline phosphatase positive colonies, normalized to the 0 hr sample (right). *n* = 5 biologically independent replicates. AP, alkaline phosphatase. Values in **f,h,i** are means ± SEM; Two-sided Student’s t-tests were used for the two conditions. *P* Values are indicated above each time point. Source numerical data and unprocessed blots are available in Source data.

Adding both auxin and dTAG13 leads to a uniform and almost complete depletion of SUV39H1 and SETDB1 proteins within 1 hour (**Fig. 1b and Extended Data Fig. 1f-g**). Western blot analysis of chromatin fractions reveals a rapid loss of H3K9me3, with ∼55% loss of H3K9me3 in chromatin within 12 hr and ∼90% loss at 24 hr of the KMTs’ degradation (**Fig. 1b and Extended Data Fig. 2a**). Mass spectrometry analysis of acid-extracted histones from nuclei shows that H3K9me3 is the most depleted histone mark at 48 hr of degradation, while the repressive heterochromatin mark H3K27me3 is largely unaltered and H3K9me2 is modestly elevated within 48 hr (**Fig. 1c,d and Extended Data Fig. 2b-d**). Interestingly, the H4K20me3, also associated with heterochromatin^30^ was mostly unaltered within 12 hr but reduced by ∼50% at 24 hr and by ∼80% at 48 hr (**Extended Data Fig. 2b-d**). The slower H4K20me3 decay indicates its dependency on the H3K9me3 machinery. Mass spectrometry analysis of total nuclear H3K9me3, which would include histones not yet integrated into chromatin^23^, shows faster decay kinetics than the chromatin fraction, with ∼90% of total H3K9me3 loss within 12 hr, and almost complete depletion within 24 hours (**Fig. 1d**). The rapid H3K9me3 loss enables us to identify the primary consequences of KMT and H3K9me3 depletion, compared to conventional genetic deletion approaches^4,25,26^. Considering a ∼17 hr doubling time of dTKO mESCs (**Extended Data Fig. 2e**), the results indicate that H3K9me3 decay is faster than expected from passive replicative dilution alone.

We found that 6 hr of thymidine treatment blocks ESC DNA synthesis by 99.9%, effectively eliminating passive replicative dilution (**Extended Data Fig. 3a and Supplementary Fig. 3a-c**). Such treatment, coupled with dual degron application, extends the half-life of H3K9me3 in chromatin from ∼11 to ∼20 hr (**Fig. 1e,f and Extended Data Fig. 3b**). Thus, passive dilution is a major contributor to H3K9me3 decay when the relevant KMTs are degraded. However, after prolonged depletion of KMTs for 24 hours, H3K9me3 levels are eventually reduced to a level similar to cells without thymidine treatment (**Fig. 1e,f**), indicating that active removal mechanisms also contribute to the H3K9me3 decay.

To assess the basis for active removal, we focused on H3K9me3 demethylases KDM4A- C^31–33^. Knocking down KDM4A-C simultaneously or chemically inhibiting KDM4 activities (**Extended Data Fig. 3c-e**) delays the half-life of H3K9me3 decay from ∼11 hr to ∼19 hr (**Fig. 1g,h and Extended Data Fig. 3b**). Thus, both passive replicative dilution and active removal by KDM4s contribute to H3K9me3 dynamics in pluripotent mammalian cells.

The AID and dTAG degradation systems are partially reversible^27,28^. After 12 hr of KMTs’ degradation, followed by 24 hr of dual degron washout, despite a rapid recovery of SUV39H1 and, to a lesser extent, of SETDB1, global H3K9me3 is not recovered to the original level (**Extended Data Fig. 3f-g**). Thus, restoring H3K9me3 levels by the “reader-writer” self-enforcing program requires a threshold level of H3K9me3 mark and KMTs. Consequently, more than 60% of the degron-treated cells irreversibly lost their self-renewal capacity after 12 hr and the remainder was eliminated after degradation for 48 hr (**Fig. 1i,j**), whereas dual degron treatment of the parental untagged cTKO mouse ESCs had no effect on self-renewal (**Extended Data Fig. 3h**). Indeed, within 48 hr of degradation, IAP, Major Satellite, LINE1 and MERVL repeats, and totipotent gene Zscan4 are activated, whereas the pluripotent genes Oct4 and Nanog are downregulated and early differentiation markers are not activated (**Extended Data Fig. 4a**). Therefore, maintaining pluripotency requires continuous H3K9me3 KMT activities to counteract constant erosion of H3K9me3 by passive and active removal mechanisms.

Extended treatment of degrons for 2-7 days led to activation of various lineage markers, including FoxA2, T/brachyury, Cdx2, and GATA6, and more flattened and enlarged cell shapes, clear indications of ES cell differentiation (**Extended Data Fig. 4a-b**), showing how H3K9me3 heterochromatin suppresses diverse lineage programs. Prolonged H3K9me3 loss for more than 2 days also led to increased nuclear shape irregularities, an aberrant DNA content profile, and growth arrest (**Extended Data Fig. 2e, Extended Data Fig. 4c-d**). Therefore, rapid loss of H3K9me3 within 48 hr leads to a cascade of events culminating in the dissolution of the pluripotency network. To investigate the primary effects of KMTs and H3K9me3 loss, we focused on the first 48 hr of degradation in the following studies.

### Chromatin HP1 engagement requires both KMTs and H3K9me3

To investigate the kinetics of H3K9me3-heterochromatin loss and its impact on chromatin structure and gene expression, we performed spike-in normalized ChIP-seq^34^ for H3K9me3 and HP1β, ATAC-seq, nascent RNA TT-seq (**Fig. 2a, and Supplementary Fig. 4a-e**), and mass spectrometry of the chromatin fraction to investigate how H3K9me3 loss affect H3K9me3- associated proteins’ binding to chromatin upon KMTs’ degradation (**Extended Data Fig. 5a**).

**Fig. 2:**
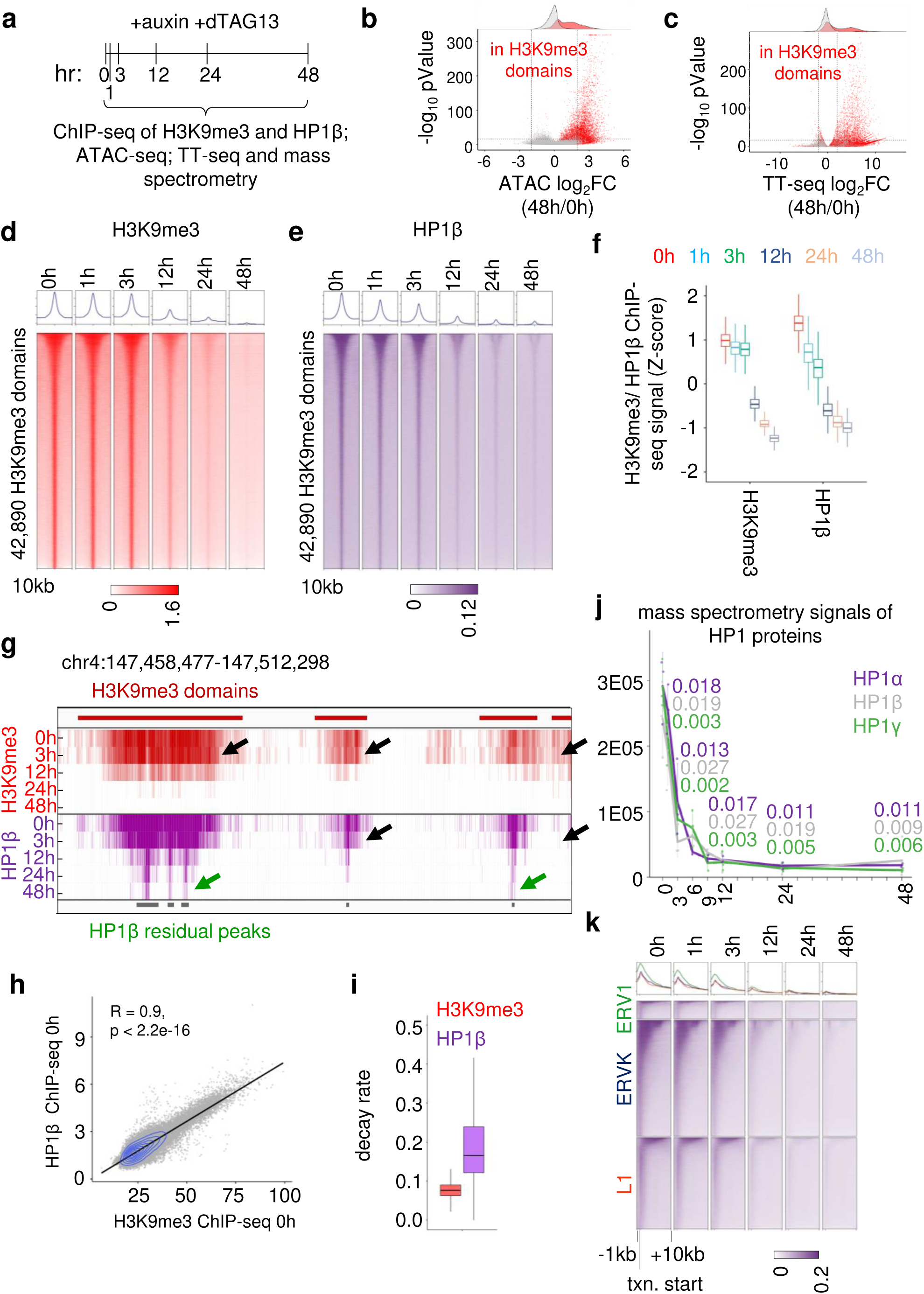
H3K9me3 and HP1β decay rates differ genome-wide **a**, Schematics of H3K9me3-KMTs’ degradation time course for H3K9me3 and HP1β ChIP-seq, ATAC-seq and TT-seq, and mass spectrometry analysis (see Methods section for details). **b,c,** Differential analysis of the ATAC-seq (**b**) and nascent RNA TT-seq (**c**) between 48 hr and 0 hr of degradation. The ATAC peaks that directly overlap with the H3K9me3 domains (**b**) or the transcription units within 10kb of the nearest H3K9me3 domains (**c**) are highlighted in red. **d,e,** Heatmaps of H3K9me3 (**d**) and HP1β (**e**) ChIP-seq signals at 42,890 H3K9me3 domains over the time course. Domains were ranked in descending order of their size and centered at the middle of each domain with 10 kb flanking windows. **f**, Boxplots comparing H3K9me3 and HP1β ChIP-seq signals (Z-score transformed) at 42,890 H3K9me3 domains over the degron time course. **g**, Genomic snapshot of H3K9me3 (red) and HP1β (purple) ChIP-seq tracks over the degradation time course. The genome coordinate (mm10) is indicated above. The black arrows highlight faster HP1β loss than the H3K9me3 mark at the corresponding domains. The green arrows indicate the residual HP1β peaks after complete H3K9me3 depletions. **h**, Scatter plot of H3K9me3 and HP1β ChIP-seq signals (spike-in normalized ChIP/Input) at H3K9me3 domains in dTKO cells before degradation. **i**, Boxplot of H3K9me3 and HP1β decay rate across the H3K9me3 domains (n = 42,890). **j**, Mass spectrometry quantifications of the HP1α/β/γ in the chromatin fractions following KMTs’ degradation. Values are means ± SEM.; *n* = 3 biologically independent replicates. Two-sided Student’s t-tests were used for each time point vs. 0 hr. *P* Values of HP1α/β/γ at 3, 6, 12, 24 and 48 hr times are indicated above the graph. **k**, Heatmaps of HP1β signals over the degradation time course at ERV1, ERVK and L1 repeats that intersect with 24 hr residual HP1β peaks. ERV, endogenous retroviral elements; L1, LINE1. In boxplots (**f,i**), the boxes represent interquartile range (IQR) from 25th to 75th percentile, with the median at the center, and whiskers extending to 1.5 times the IQR from the quartiles. Source numerical data are available in Source data.

Based on H3K9me3 changes measured by Western blotting (**Fig. 1b**), we selected following times for genomic analysis: 1 hr and 3 hr, when KMTs are depleted but chromatin-bound H3K9me3 is largely unchanged, 12 hr, when H3K9me3 is about half-way depleted, and 24 and 48 hr, when H3K9me3 is mostly depleted. As expected, by 48 hr we observed more chromatin and transcriptional activation responses than repression, most of which occurred within H3K9me3 domains (**Fig. 2b,c**, red dots). The findings illustrate the specificity in phenotype obtained by the degron approach and contrast with the secondary gene repression responses seen with the slower genetic deletions, which take days to deplete the KMTs^4,25,26^.

Broad H3K9me3 domains (42,890 in total) were called using RSEG^35^ and H3K9me3 and HP1β signals within the domains were quantified over the time course. Consistent with the Western blot data after dual degron treatments, the average genomic H3K9me3 signals of all the H3K9me3 domains was reduced by ∼16% within 3 hr, by ∼50% at 12 hr and by ∼80% at 24 hr (**Fig. 2d-f,g** upper arrow). In control ESCs, HP1β and H3K9me3 levels highly correlate across H3K9me3 domains (**Fig. 2d-e,g,h**). Upon degron treatment in dTKO cells, at the genomic heterochromatin domain level, HP1 is reduced ∼40% by 3 hr, and ∼95% by 12 hr (**Fig. 2d-g**). Fitting the kinetics of the H3K9me3 and HP1β/γ loss at each H3K9me3 domain with an exponential decay model, we found that decay rates of HP1β are indeed higher than for the H3K9me3 mark across H3K9me3-domains over the 48 hr time course (**Fig. 2i**). Considering a lack of suitable antibodies for HP1α and HP1γ ChIP-seq, we performed mass spectrometry and Western blotting of the chromatin fraction to discern the three HP1 isoforms and found similar dissociation kinetics among bulk HP1α/β/γ from chromatin, with all three HP1 isoforms dissociated as early as 3 hr (**Fig. 2j and Extended Data. Fig. 5b**), Yet notably the total cellular level of HP1α/β/γ protein is largely unchanged within 48 hr (**Extended Data Fig. 5c**). Thus, stable HP1 binding across H3K9me3 domains requires HP1-KMT interactions, in addition to recognizing H3K9me3 marks, as indicated by previous in vitro biochemical assays^13^.

Unexpectedly, although H3K9me3 signals are largely depleted by 24 hr of degron treatment, 8712 residual peaks for HP1β (median size of 240 bp) remained and 5917 peaks persist at 48 hr (**Fig. 2g, green arrows**), 87% of which fall within the starting H3Kime3 domains (median size of 3 kb, **Extended Data Fig. 5d**). The residual peaks primarily mark LTR and promoter regions of ERV1, ERVK, and LINE1 repeats (**Fig. 2k and Extended Data** Fig. 5e) and are enriched for TRIM28, MPP8, METTL3, and the m^6^A reader YTHDC1 in normal ESCs (**Extended Data** Fig. 5f); these proteins were shown to recruit KMTs to chromatin in mESCs^36,37^. Notably, the residual HP1β peaks are largely devoid of the H3K9me2 mark (**Extended Data** Fig. 5f) that, in principle, can also be recognized by HP1^38^. Therefore, receding of H3K9me3 upon KMTs’ degradation unexpectedly reveals HP1 at residual sites characteristic of H3K9me3- heterochromatin nucleation.

ChIP-qPCR of TRIM28/KAP1, METTL3, and YTDC1 confirmed their binding at six HP1 residual peaks before degradation (**Extended Data** Fig. 5g-j). Upon KMTs’ depletion, METTL3 and YTHDC1 binding at all six HP1 residual peak regions were reduced by ∼50% at 12 hr, and were mostly depleted at 48 hr, kinetics similar to H3K9me3 decay (**Extended Data** Fig. 5h-i). In contrast, TRIM28/KAP1 binding was largely stable in five of the six residual peak regions, and was reduced by ∼ 50% at 48 hr (**Extended Data** Fig. 5j). Thus, TRIM28/KAP1 remains after H3K9me3 domain decay and may maintain the residual HP1 binding on chromatin, while H3K9me3 and other heterochromatin associated proteins are lost.

Furthermore, consistent with the Western blot data, H3K27me3 ChIP-seq signals during the 48 hr time course are mostly unaltered (**Extended Data** Fig. 6a-b). The H3K9me2 signals are slightly elevated at both the lost H3K9me3 domains and the H3K9me2 domains (**Extended Data** Fig. 6c-d), indicating a weak compensation of H3K9me3 loss by H3K9me2. In comparison, H4K20me3 levels correlate well with H3K9me3 at the H3K9me3 domains before degradation, are largely stable within 12 hr of degradation, and were reduced by ∼50% at 24 hr and reduced by ∼80% at 48 hr across the H3K9me3 domains (**Extended Data** Fig. 6e-f), further confirming the dependencies of H4K30me3 mark on H3K9me3 machineries across the genome.

### H3K9me3 heterochromatin domains have four stability types

To better understand the functional diversities of H3K9me3 domains, based on the relative timing of H3K9me3 loss across the genome, we first partitioned H3K9me3 domains into four clusters (see alluvial plots in **Fig. 3a**): an early-cluster that lost H3K9me3 by 3 hr (1,498 domains), an intermediate-cluster by 12 hr (30,660 domains), a late-cluster by 24 hr (8,772 domains), and a residual-cluster (1,960 domains) that remains after 24 hr. The early-, intermediate-, and late- cluster have low, intermediate, and high initial H3K9me3 and HP1β signals, respectively, whereas the residual H3K9me3 domains paradoxically have an intermediate H3K9me3 level, but minimal HP1β, suggesting a distinct chromatin state, as detailed below (**Fig. 3b and Extended Data** Fig. 7a-b). After fitting the kinetics of H3K9me3 decay with exponential decay models, the decay rates of H3K9me3 at early, intermediate, and late clusters are incrementally lower (**Fig. 3c**), indicating that different heterochromatin clusters vary in their stabilities and that timing and kinetics of H3K9me3 loss are largely proportional to their initial H3K9me3 and HP1 signals. Most significantly, the four H3K9me3 stability clusters harbor distinct transcription factor motifs and mark different genetic pathways and DNA repeats (**Extended Data** Fig. 7c-e). For example, the early- and residual- clusters of H3K9me3 domains contain fewer DNA repeats than the intermediate- and late- H3K9me3 domains (**Extended Data** Fig. 7e). Thus, the alluvial H3K9me3 clusters associated with different initial H3K9me3 levels and stabilities resolve distinct functional H3K9me3 domains.

**Fig. 3:**
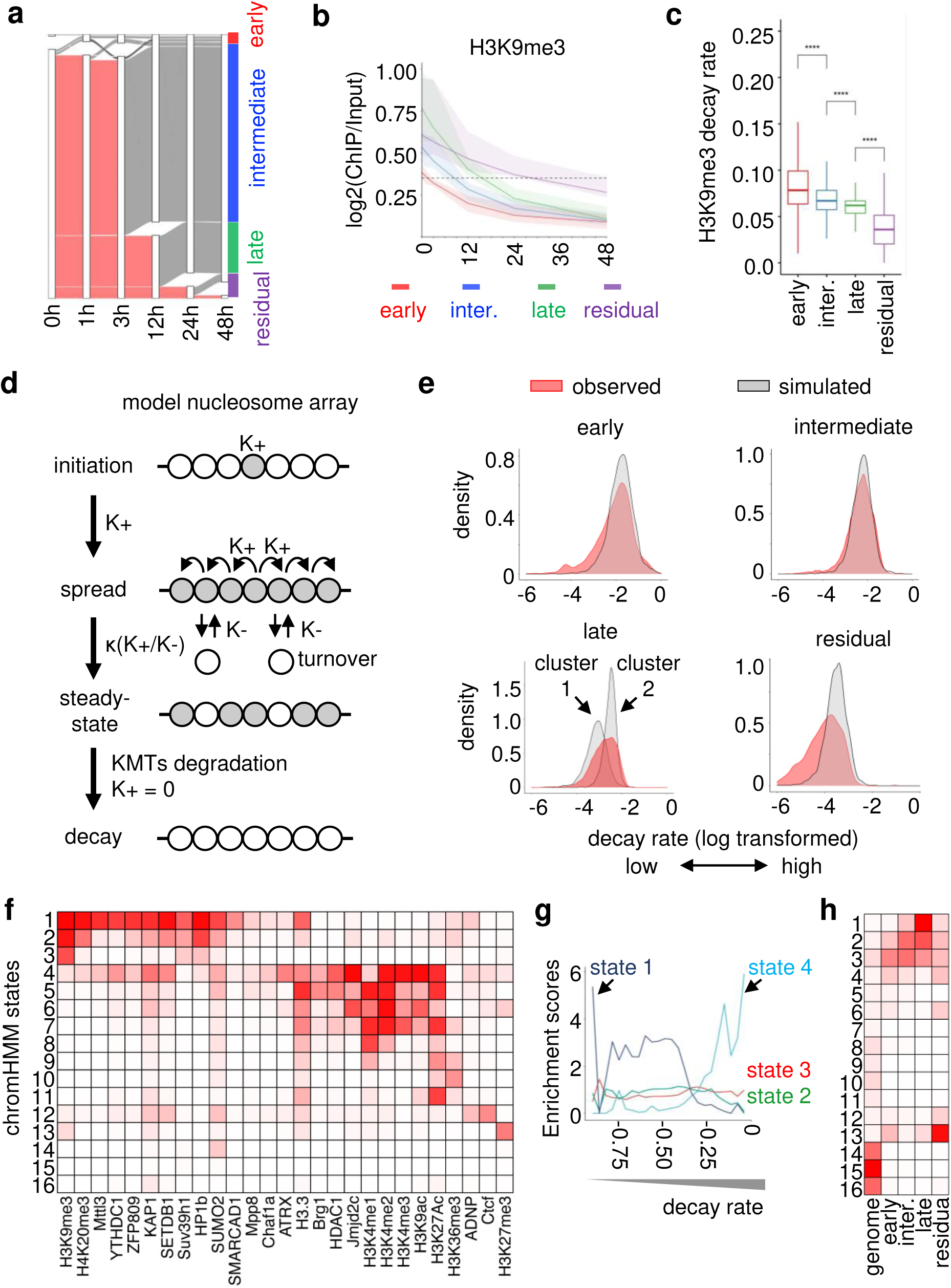
Heterochromatin states influence kinetics of H3K9me3 decay **a**, Alluvial plots showing different timings of the H3K9me3 loss. The H3K9me3 of early-, intermediate-, and late- cluster are lost at 3 hr, 12 hr, and 24 hr, respectively, whereas the residual- cluster persist after 24 hr. Grey ribbons indicate the loss of the H3K9me3, and the height of the ribbon is proportional to the number of domains. **b**, Ribbon plot showing the different kinetics of H3K9me3 decay at each H3K9me3 alluvial cluster. The ribbon represents the standard deviations of the H3K9me3 signals from the mean value at each cluster. **c**, Boxplots showing the distributions of H3K9me3 decay constant at different H3K9me3 clusters: early (n=1498), intermediate (n=30660), late (n=8772), and residual (n=1960). The boxes represent interquartile range (IQR) from 25th to 75th percentile, with the median at the center, and whiskers extending to 1.5 times the IQR from the quartiles. Statistical analysis was performed using Wilcoxon test. *P* values are 7.1E-86, 1.8E-237 and 3.2E-234. ****, P < 2.22e-16 **d**, Schematics of mathematical modeling of the H3K9me3 establishment and decay. The K+ is the aggregated rates of the nucleation and spreading of H3K9me3 mark; K- is the aggregated decay rates eroding the H3K9me3. **e**, Density plots comparing the distributions of observed H3K9me3 decay constant (red) at different H3K9me3 clusters with the simulated H3K9me3 decay constant determined by different K- (grey). Note that there are two sub-clusters in the slow- cluster fitted with different K- parameters. **f**, Heatmap of the 16-state ChromHMM model parameters (state numbers on the left). Columns indicate the enrichments of different chromatin features at each chromatin state. Note that H3K9me3-heterochromatin (first column) are partitioned in chromatin states 1-4. **g**, Jaccard enrichment scores of heterochromatin states 1-4 at H3K9me3 domains with different ranges of decay rates. **h,** Heatmap showing the enrichments of each H3K9me3 clusters at different chromatin states. The percentage of each state in the mouse genome is indicated in the first column. Source numerical data are available in Source data.

To understand principles governing different stabilities of H3K9me3 domains, we applied a mathematic model that simulates the H3K9me3 level at a given domain using the ratio (κ) of K+, the aggregated rates of H3K9me3 nucleation and spread, and K-, the aggregated rate of H3K9me3 turnover (κ = K+/K-)^19,24^ (**Fig. 3d and Supplementary Fig. 5a,b**). Different from previous models that assume same parameters for all domains^24^, given our observations of different stability types, we swept both K+ and K- parameters (hence, different κ ratios) in the model to fit both initial H3K9me3 levels and decay rates observed at different H3K9me3 clusters (**Supplementary Fig. 5c-d**). We found that K- parameters in the model, dictating the decay rate of the H3K9me3 signals are progressively lower at the early, intermediate, late and residual clusters (**Fig. 3e**), and the decay rate of the late H3K9me3 cluster shows a bimodal distribution, despite a similar overall κ (K+/K-) (**Fig. 3e**), indicating modulators of H3K9me3 levels and domain stability.

### Different chromatin states influence H3K9me3 stability types

To elucidate how the local chromatin environment modulates H3K9me3 domain stabilities, we developed a 16-state ChromHMM model from ESC data^39^ that reveals 4 different H3K9me3 heterochromatin states with distinct chromatin features (**Fig. 3f**, states 1-4). State 1 has highest enrichment for the heterochromatin nucleation complexes, such as KMTs, ZFP809, TRIM28 (**Figure 3f and Extended Data** Fig. 8a), and correlates with the earliest H3K9me3 domains during pre-implantation development^40^ (**Extended Data** Fig. 8b). Consistent with the HMM model, HP1 residual peaks at 48 hr are highly enriched in state 1 (**Extended Data** Fig. 8c). State 2 and state 3 often flank state 1 within the same initial, pre-degron H3K9me3 domain and apparently mark the spreading and edges of H3K9me3 domains respectively, with decreasing KMTs, HP1, and H3K9me3 levels (**Extended Data** Fig. 8a,d**,e**). State 4 has lower H3K9me3 and HP1, but is characterized by co-occurring heterochromatin and euchromatin features, such as KDM4, H3.3, and MPP8, a component of HUSH complex^20^ (**Fig. 3f and Extended Data** Fig. 8a). Therefore, our ChromHMM model captures diverse functional heterochromatin states.

To understand how different heterochromatin states influence H3K9me3 domain stabilities, we grouped H3K9me3 domains into bins with increasingly higher decay rates (**Fig. 3g**, left to right on x-axis) and found that HMM-based heterochromatin states 1 and 4 are associated with the lowest and highest decay rates, respectively (**Fig. 3g**). The early-lost cluster is more enriched in chromHMM-heterochromatin states 3 and 4, whereas intermediate- and late-lost clusters are increasingly more enriched in heterochromatin state 1 (**Fig. 3h**). The high enrichment of KDM4, H2A.Z, H3.3 and its chaperone ATRX, and chromatin remodeler Brg1 in heterochromatin state 4 (**Fig 3f and Extended Data** Fig. 8a) are consistent with active demethylation and histone turnover contributing to the high decay rates (K-) in the early-lost domains (**Fig 3e**). Notably, HDACs, also enriched in state 4, suppress histone turnover^41^ and our mass spectrometry analysis shows a rapid dissociation of HDAC1/2/3 from the chromatin upon KMTs’ degradation, potentially exacerbating H3K9me3 loss by increased histone turnover (**Extended Data** Fig. 8f). State 4 predominantly marks developmental genes and LINE and SINE elements, especially evolutionary young LINE1 subfamily L1Md_T (**Extended Data** Fig. 8g).

In contrast to the rapid loss of HDACs after dual degron treatment, our mass spectrometry analysis shows that many H3K9me3-associated proteins enriched in the chromHMM- heterochromatin state 1, including SUMO, TRIM28, and ZFPs remain on the chromatin during H3K9me3 loss (**Extended Data** Fig. 8f). Their more stable association with H3K9me3 domains at intermediate and late-clusters would counteract rapid erosions of H3K9me3 domains upon the KMTs’ degradation (**Fig. 3c**). State 1 mainly marks LTR retrotransposons, especially IAPs (ERVK subfamily) (**Extended Data** Fig. 8g). The residual cluster is primarily enriched in state 13, with high H3K27me3 but low H3K9me3 signals, and marks developmental genes regulated by Polycomb (**Fig 3f,h**). We previously observed such domains with H3K9me3 and H3K27me3 in somatic cells^21,22^, and recent mass spectrometry analysis of dual H3K9me3 and H3K27me3 modifications shows enrichment for non-canonical PRC1 complex reader proteins in HMM state 13^42^. Accordingly, mass spectrometry analysis shows that Suz12, a PRC2 subunit, is drastically increased in chromatin within 3 hr of KMTs’ loss, with a coordinate elevation of H3K27me1, but not the other core PRC2 components Eed or Rbbp4 or H3K27me3 levels (**Fig. 1c and Extended Data** Fig. 8f). We conclude that the PRC2 proteins quickly sense H3K9me3 loss, especially at the developmental genes, which could counteract rapid H3K9me3 erosion. Thus, our acute KMT degron approach with mathematic modeling, mass spectrometry, and global genomic analyses reveals that H3K9me3 stability at different genetic networks and repeat families is not monotonic across the genome. Different local chromatin environments predict the four H3K9me3- heterochromatin stability types.

### A threshold level of HP1 governs heterochromatin integrity

Next, we assessed how different stabilities of H3K9me3 domains impact the chromatin structure and expression of resident genes and regulatory elements. The scatter plots of H3K9me3 or HP1 vs. ATAC signals, as a measure of chromatin accessibility, show clear anti-correlations (**Fig. 4a,b**, at 0 hr), consistent with HP1 oligomerization and multivalent interactions with H3K9me3-marked nucleosomes compacting H3K9me3-heterochromatin^43^. During dual degron treatment, an increase in chromatin accessibility starts as early as 3 hr, concomitant with the rapid dissociation of HP1 from the H3K9me3-heterochromatin (**Fig. 4a-c and Extended Data** Fig. 9a). Significantly, the increase of ATAC signals were observed when HP1 and H3K9me3 are reduced, but not fully diminished (**Fig. 4a,b**, compare dots above red lines at 3 hr), indicating that after KMTs’ depletion, maintaining heterochromatin compaction is sensitive to a threshold level of H3K9me3 and HP1, below which heterochromatin compaction is lost. With the initial level of HP1 at early-, intermediate-, and late- domains increasingly higher (**Fig. 4d,e**), the initial ATAC signals are progressively lower (**Fig. 4c and Extended Data** Fig. 9b) and the timing and kinetics of chromatin opening (**Fig. 4f**) and transcriptional activation (**Fig. 4g**) are correspondingly slower. Therefore, the differences in kinetics of HP1 loss at the four domain stability types, proportional to the initial H3K9me3 and HP1 levels (**Fig. 4d,e**), predicts the different timings of chromatin opening and target gene activation after the degron treatment. Notably, the bulk diminution of H3K9me3 (**Fig. 4d**) occurs concomitant with or after the increase of chromatin opening and transcription at H3K9me3 domains (**Fig. 4d-g**), indicating how a partial, critical loss of heterochromatin integrity, prominently at 12 hr, leads to rapid transcriptional de-repression.

**Fig. 4:**
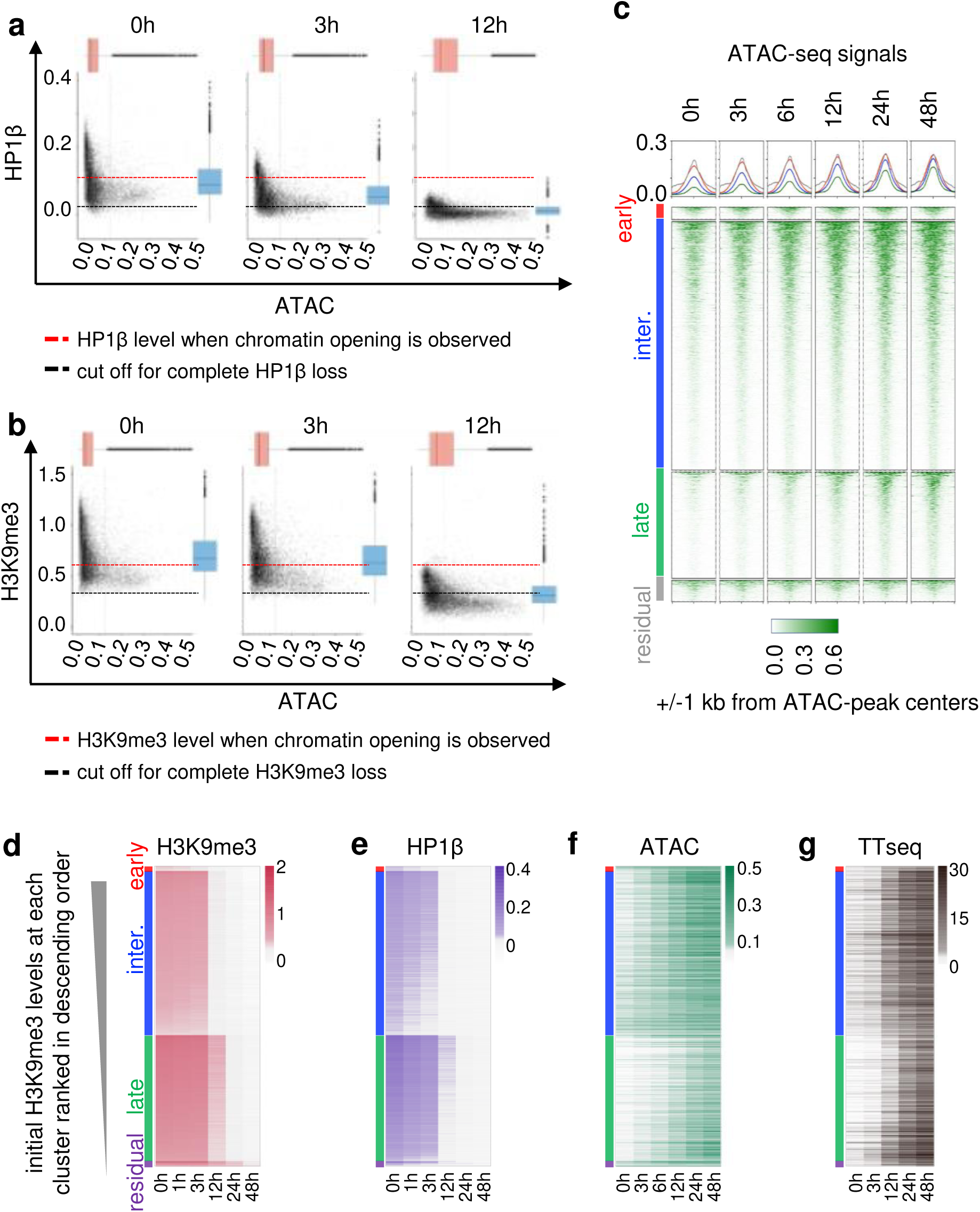
Maintaining heterochromatin integrity involves a threshold level of H3K9me3 and HP1 **a,b,** Scatter plots showing the signals of HP1β (**a**) and H3K9me3 (**b**) vs. ATAC peaks within the H3K9me3 domains at 0 hr, 3 hr, and 12 hr of degradation. Boxplots on the top and on the right of the scatter plots are the distributions of ATAC signals (n=16783) and HP1β signals (**a,** n=12505) or H3K9me3 (**b**, n=12505), respectively. The black dashed horizontal lines indicate the threshold of complete loss of HP1β (**a**) or H3K9me3 (**b**). The red dashed horizontal lines indicate the threshold of HP1β (**a**) or H3K9me3 (**b**) signals at which ATAC signals were observed. **c**, Heatmaps showing ATAC signals within 1 kb window flanking the center of ATAC peaks at different H3K9me3 clusters over the degradation time course. **d-g,** Supervised heatmaps showing higher H3K9me3 (**d**) and HP1β (**e**) signals results in slower kinetics of chromatin opening in ATAC-seq (**f**) and transcription activation in TT-seq (**g**) at each H3K9me3 domains at different H3K9me3 clusters over the time course. The H3K9me3 domains within each cluster were ranked in descending order based on H3K9me3 signals at 0h (**d**), and the same order was applied to the HP1β (**e**), and ATAC peaks within the H3K9me3 (**f**) and the nearest transcription units in TT-seq (**g**). Source numerical data are available in Source data.

Indeed, over the dual degron time course, we observed that 1055 protein coding genes are activated, with immune responsive genes, male meiosis-related genes, and the master regulator of totipotency Zscan4 being activated as early as 3 hr (**Fig. 5a-b**). We conclude that these developmental genetic programs are tightly linked to heterochromatin dynamics. Furthermore, 299 (28%) of the 1055 activated genes are involved in diverse cell lineages (**Fig. 5b**) and are significantly activated within 12 hr of degradation. Concomitantly, 8309 transposable elements are activated over the time course, in which 3134 (38%) were activated within 12 hr (**Fig. 5c**). We observed that 36% of activated protein coding genes have an activated transposable element within 5 kb of their transcription start site (**Fig. 5d**), suggesting a coordinated activation of H3K9me3-repressed transposable elements and nearby protein coding genes. Furthermore, 45% of the genes and 43% of the DNA repeats that are de-repressed by 12 hr of KMTs’ degradation are also activated in HP1α/β/γ triple knock-out mouse ESCs^44^ (**Fig. 5e,f**), consistent with the notion that rapid dissociation of HP1 proteins causes loss of heterochromatin integrity. Notably, H3K27me1 is normally observed at actively transcribed genes^45^ and the increase of the H3K27me1 at 12 hr (**Extended Data** Fig. 2b) coincides with global transcriptional de-repression in H3K9me3 domains. The critical loss of heterochromatin integrity within 12 hr leading to activation of various lineage genes and transposable elements explains the time point’s irreversible exit from pluripotency in mouse ES cells (**Fig. 1i-j**).

### Pioneer factors elicit further H3K9me3 loss after KMTs’ decay

To investigate when and which transcription factors can access the newly de-compacted H3K9me3-heterochromatin to induce heterochromatin remodeling and cell fate transitions during development, we surveyed footprints of all the expressed transcription factors in mESCs within ATAC peaks^46^ near activated genes and DNA repeats. We included an intermediate 6 hr time point in the ATAC-seq experiments to investigate the transcription factor actions at a finer level. ATAC footprinting^46^ indicates that, after KMTs’ depletion, numerous transcription factors access H3K9me3 domains at the activated genes and DNA repeats, concomitant with the rapid HP1 loss from 3 hr to 12 hr (**Fig. 6a and Extended Data** Fig. 9c). We suspected that pioneer transcription factors that target nucleosomal DNA could be involved in gene activation upon the KMTs’ degradation, because mass spectrometry did not reveal an overt loss of core histones at early time points (**Extended Data** Fig. 8f). Indeed, footprints for pioneer factors NFYA, POU5F1 (OCT4), and KLF4 appear in heterochromatin within 3 hr (**Fig. 6b,c and Extended Data** Fig. 9d) and only later generate nucleosome-free regions at their footprinting sites, characteristic of euchromatin (**Fig. 6d**), which further leads to chromatin opening and transcriptional activation (**Fig. 6e-g**). This is consistent with mass spectrometry data showing later binding of chromatin remodelers SMARCAD1, SMARCA2 and CHD1 during the degron time course (**Extended Data** Fig. 8f).

**Fig. 5:**
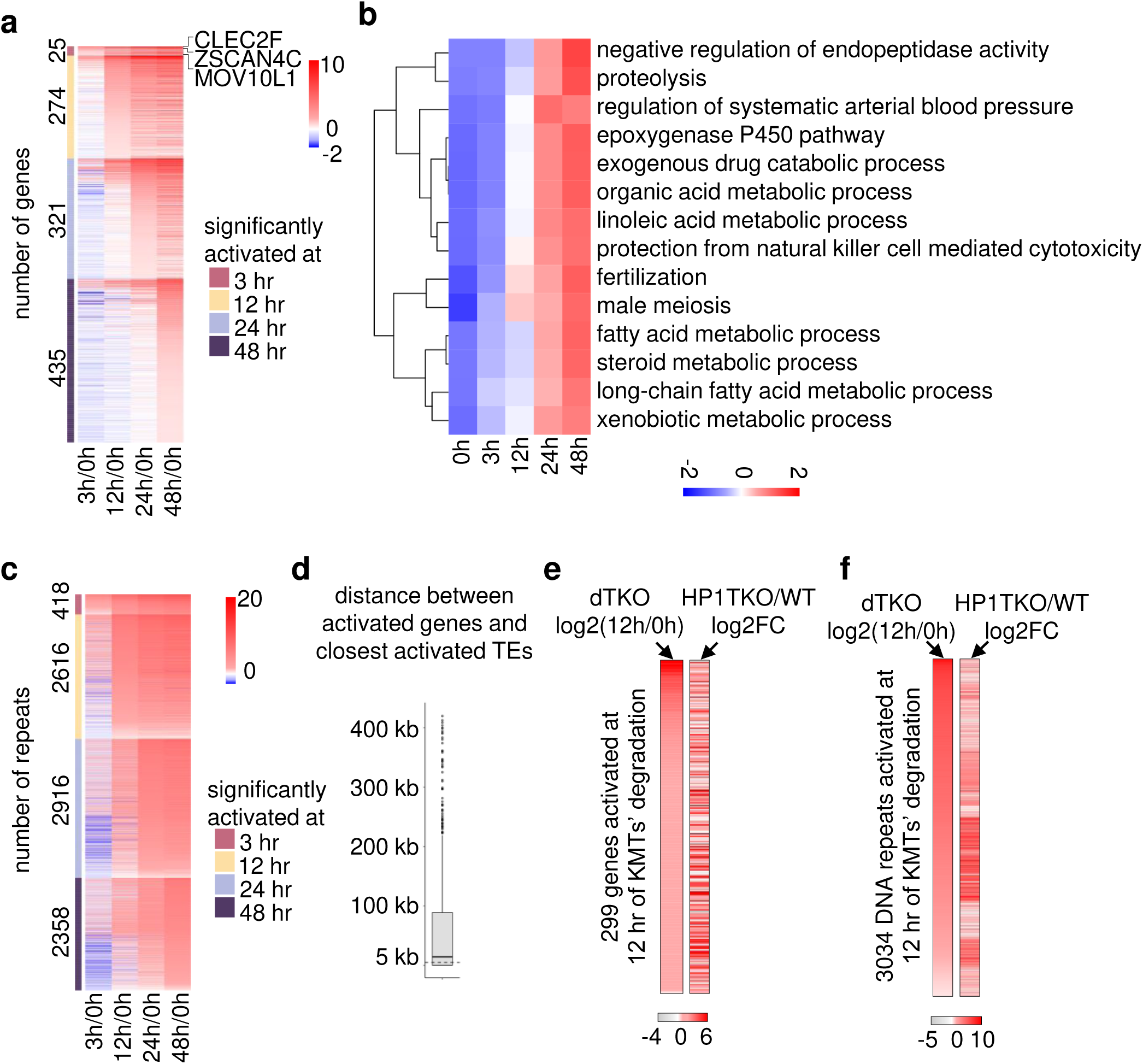
The critical loss of heterochromatin integrity within 12 hr leads to activation of various lineage genes and DNA repeats **a**, Heatmaps showing 1055 genes that are significantly activated at different time of KMTs degradation. The values are log2 fold changes of averaged expression level at each time point normalized to 0 hr. **b**, Heatmap showing the transcriptional changes of different GO functional terms activated over the time course. The values are averages of genes within each GO terms after Z-score transformation. **c**, Heatmaps showing 8309 transposable elements that are significantly activated at different time of KMTs degradation. The values are log2 fold changes of averaged expression level at each time-point normalized to 0 hr. **d**, Boxplot showing the distances between activated genes (n=1055) and the nearest activated transposable elements. The boxes represent interquartile range (IQR) from 25th to 75th percentile, with the median at the center, and whiskers extending to 1.5 times the IQR from the quartiles. **e**, Supervised heatmaps showing on the left, the log2 fold change of 299 genes significantly activated at 12 hr of KMTs degradation (in descending order). The same gene order was applied to heatmap on the right showing the log2 ratio of HP1 TKO/WT mouse ESCs (GSE210606). **f**, Supervised heatmaps showing on the left, the log2 fold change (in descending order) of 2100 transposable elements significantly activated at 12 hr of KMTs degradation that are also detectable in the published HP1 TKO RNA- seq data (GSE210606). The same order was applied to heatmap on the right showing the log2 ratio of HP1 TKO/WT mouse ESCs. Source numerical data are available in Source data.

**Fig. 6:**
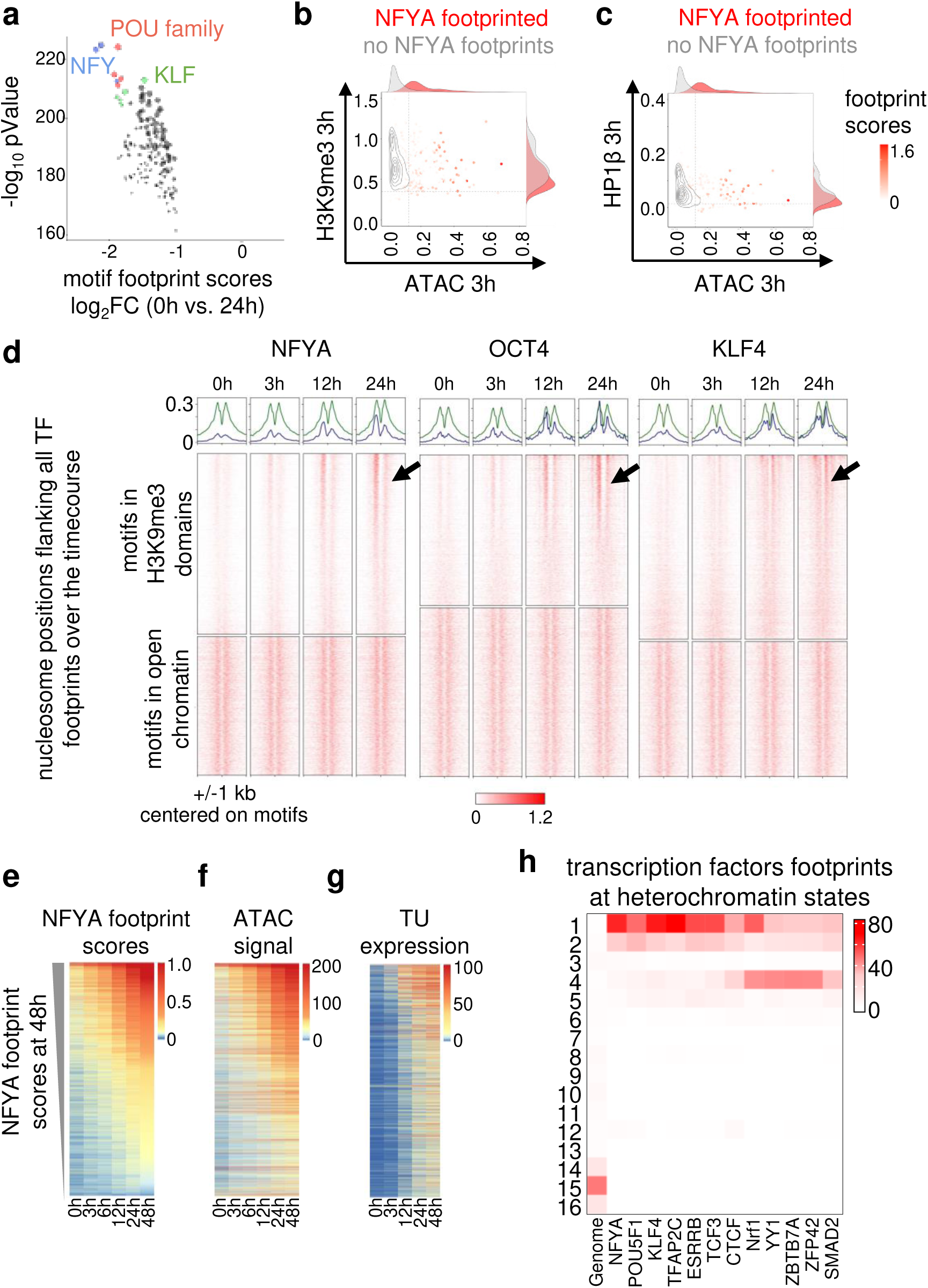
Pioneer factors eliciting further heterochromatin loss after KMTs’ decay **a**, Differential analysis of the footprint scores of the transcription factors expressed in mouse ESCs between 24 hr and 0 hr of degradation. The NFYA/B/C, POU, and KLF transcription factors are highlighted. **b,c,** Scatter and contour plots of signals of H3K9me3 (**b**) or HP1β (**c**) in H3K9me3 domains vs. the ATAC peaks within the H3K9me3 domains at 3 hr. The NFYA footprint scores were super-imposed on the dot plots. **d**, Heatmap showing the nucleosome positioning within 1 kb window flanking the NFYA, OCT4 and KLF4 footprinted motifs over the degradation time- course. Nucleosome positions were inferred from ATAC-seq data using nucleoATAC. **e-g**, Supervised heatmaps showing NFYA footprint scores ranked based on footprint scores at 48 hr in descending order (**e**), and the associated ATAC signals (**f**), and the expression of nearest transcriptional units (**g**) with the same order over the time course. **h**, Heatmap showing enrichments of transcription factors-footprinted ATAC peaks in H3K9me3 domains at different ChromHMM states. The percentage of each state in the mouse genome is indicated in the first column. Source numerical data are available in Source data.

We found low but detectable Oct4 signals by ChIP-qPCR at four predicted Oct4 footprint sites even before KMTs’ degradation, suggesting that Oct4 may sample the H3K9me3 heterochromatin in wild type cells, and Oct4 signals are increased during the degron treatment at all four sites, two of which are significantly increased within 12 hr (**Extended Data** Fig. 9e). Thus, acute KMTs’ depletion causes rapid HP1 dissociation to threshold levels, enabling pioneer factors to quickly access the newly de-compacted H3K9me3-heterochromatin, leading to chromatin remodeling and histone turnover and eliciting locally open sites.

Interestingly, the decay rates of the H3K9me3 domains with chromatin opening and transcriptional activation are higher than the decay rates of H3K9me3-domains that remain silent (**Extended Data** Fig. 9f-g). Transcription factor footprints are detected in heterochromatin state 4, associated with higher H3K9me3 decay rates (**Fig. 6h**), as well as in heterochromatin state 1, enriched in late-lost H3K9me3 domains (**Fig. 3f**). The mathematic modeling of late-lost domains revealed two sub-clusters with different decay constants (**Fig. 3e**). Notably, transcription factor footprints, including NFYA, mostly fall in the late-lost sub-cluster with a higher decay constant (**Extended Data** Fig. 9h). To test the role of NFYA in enhancing H3K9me3 decay, we performed a knock-down during the degradation time course and observed a ∼50% reduction of NFYA, which was sufficient to impair H3K9me3 loss after 12 hr of degrons’ treatment (**Fig. 7a-b**). Thus, transcription factors’ binding activates transcription at H3K9me3-heterochromatin, promoting faster H3K9me3 remodeling and decay.

**Fig. 7:**
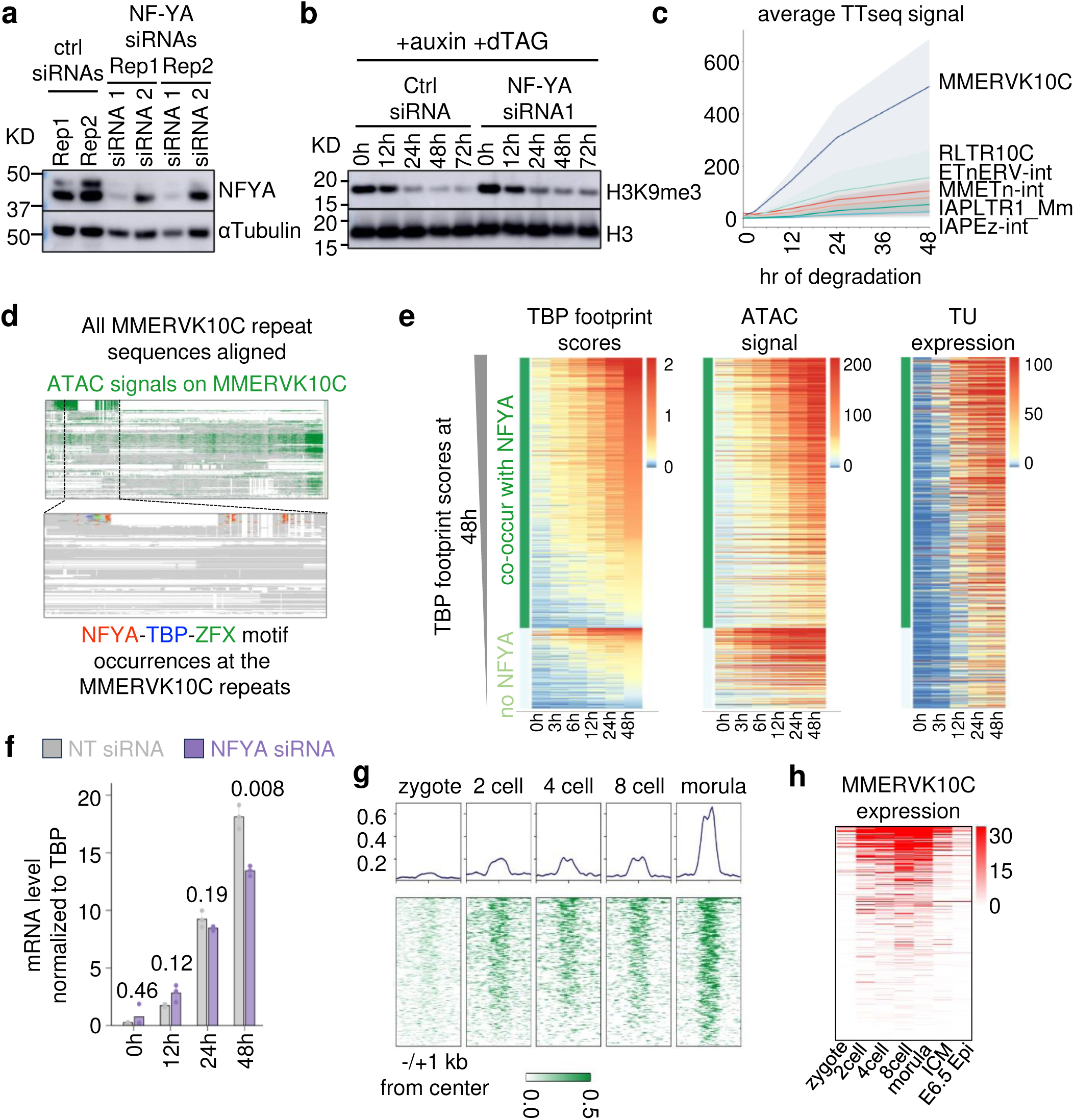
NF-YA acts as pioneer factors to uniquely target MMERVK10C repeat family **a,** Western blot of NFYA, and the loading control αTubulin after 3 days of initial siRNA transduction. **b**, Western blot of H3K9me3, and the loading control histone H3 after 3 days of initial siRNA transductions, followed by the degradation time course. The time (hr) of auxin and dTAG addition are indicated above. Experiments were repeated three times independently with similar results. The molecular weight in KD (kilodaltons) is on the left. Ctrl siRNA, scramble siRNA (**a,b**). **c,** Ribbon plots showing the expression of different repeat subfamilies over the degradation course. The lines are the means of TT-seq signal, and ribbons are the distributions of ± one standard deviation from the mean. **d,** MAFFT alignments of all MMERVK10C-int subfamily sequences (in grey, truncations in white) with ATAC signals (in green, Top) and NFYA, TBP, and ZFX motifs super-imposed on the alignments (below). **e,** Supervised heatmaps showing TBP footprint scores ranked based on footprint scores at 48 hr in descending order (**left**) with or without NFYA co-occurrences, the associated ATAC signals (**middle**), and expression of the nearest transcriptional units (**right**) with the same order over the time course. **f**, Bar graph showing RT- qPCR quantification of MMERVK10C transcript levels with NFYA footprints (normalized to housekeeping gene TBP) after 3 days of initial siRNA transductions, followed by dual degrons treatment for 0, 12, 24, and 48 hr; *n* = 3 biologically independent replicates, values are means ± SEM. Two-sided Student’s t-tests were used for the two conditions. *P* Values are indicated above each time point. **g**, Heatmaps showing changes of DNaseI-seq signals from zygote to morula stage of mouse development (GSE76642), within 1 kb window from the center of NFYA footprinted regions at MMERVK10C identified over the degradation time course. **h,** Heatmap showing expression changes of MMERVK10C with NFYA footprints over the degradation time course during early mouse development (GSE98150). Source numerical data and unprocessed blots are available in Source data.

We found that different DNA repeat families de-repressed over the degrons’ time course are associated with the appearance of particular transcription factor footprints. Most notably, NFYA and TBP footprints are highly enriched at a subset of the MMERVK10C repeats, which are activated the earliest and most robustly upon degrons’ treatment (**Fig. 7c and Extended Data** Fig. 10a). NFYA and TBP motifs are positioned closely within the activated MMERVK10C and MMERVK10C, with NFYA and TBP footprints increased more robustly than at MMERVK10Cs without NFYA footprints (**Fig. 7d-e**). Indeed, the partial knockdown of NFYA led to a partial inactivation of the MMERVK10C subfamily within 48 hr of KMTs’ degradation, compared to the control siRNA (**Fig. 7f**), confirming that NFYA activates MMERVK10C subfamily during the degrons’ time course. Furthermore, MMERVK10C subfamilies are normally activated at the 2-cell embryo stage (**Fig. 7g-h**), consistent with NFYA functioning as a pioneer factor to elicit chromatin opening and transcriptional activation in pre-implantation development^47^. Thus, different H3K9me3 stability domains restrict different transcription factors from activating lineage-specific genes and transposable elements, thereby maintaining pluripotency.

### H3K9me3-heterochromatin constrains Dppa2 to bivalent genes

We found 1,067 protein coding genes that are unexpectedly down-regulated over the degradation time course (**Fig. 8a**). Most of the down-regulated genes initially were bivalent, marked by H3K27me3 and H3K4me3 rather than by H3K9me3^48^ (**Fig. 8b**). Consistent with the genes’ down- regulation during the degron time course, H3K27me3 surrounding the genes’ transcription start sites increased (**Fig. 8c)**. Interestingly, over half (611) of the downregulated genes’ promoters are normally bound by Dppa2, which is thought to limit the PRC2 complex to maintain bivalency^49^ (**Fig. 8b**). Indeed, the H3K27me3 increase at the transcription start sites of down-regulated genes over the KMTs’ degradation time course mirrors the increase of H3K27me3 in Dppa2/4 double knock-out mESCs^49^ (**Fig. 8d**).

**Fig. 8:**
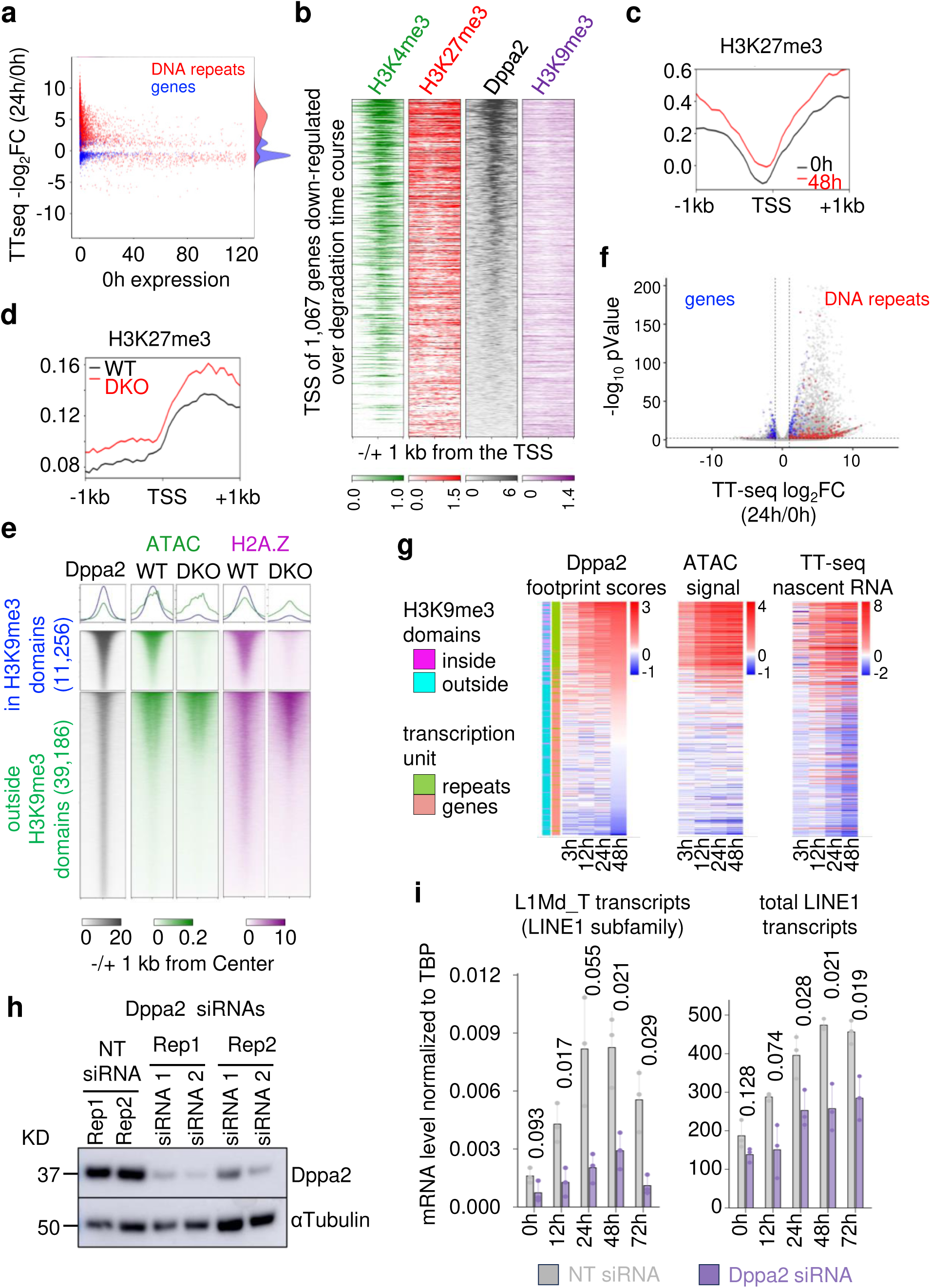
H3K9me3-heterochromatin constrains transcription factor Dppa2 to bivalent genes **a,** Differential analysis of TT-seq between 24 hr and 0 hr, highlighting DNA repeats and protein coding genes in red and blue, respectively. **b**, Heatmaps of Dppa2 (GSE117173), H3K4me3 (GSE135841), H3K27me3 (GSE135841) and H3K9me3 (this study) signals within 1kb window flanking the TSS (transcription start sites) of the down-regulated genes in the degradation timecourse. **c**, Average H3K27me3 signals within 1 kb windows flanking TSS of downregulated genes at 0 hr and 48 hr of degradation. **d,** Average H3K27me3 signals in wildtype (WT) vs. Dppa2/4 double knockout (DKO) mouse ESCs (GSE135841) within 1kb window flanking the TSS of downregulated genes in the degradation time course. **e**, Heatmaps of Dppa2 (GSE117173) in wild-type mouse ESCs, ATAC, and H2A.Z in wild-type vs. Dppa2/4 knockout (DKO) mouse ESCs (GSE135841) at Dppa2 peaks inside (top) vs. outside (bottom) the H3K9me3 domains. **f,** Volcano plot showing differentially expressed transcription units after 24 hr of degradation, with Dppa2 targets highlighted in blue (protein coding genes) and red (DNA repeats). **g**, Supervised heatmaps of log2 fold changes of Dppa2 footprint scores ranked in descending order based on the 48 hr time point (left), the associated ATAC signals (middle), and the expression of the nearest transcriptional units (right) with the same order. The annotations on the left of the heatmap indicate whether the ATAC peaks directly overlap with the H3K9me3 domains, and whether the transcription units are DNA repeats or protein coding genes. **h,** Western blot of Dppa2, and the loading control αTubulin after 3 days of siRNA transductions. Experiments were repeated three times independently with similar results. The molecular weight in KD (kilodaltons) is on the left. **e**. Bar graph comparing Dppa2 footprinted L1Md_T LINE1 subfamily transcript (left) or total LINE1 transcript levels (right) over 3 days of degradation time course with or without Dppa2 depletion. n = 3 biologically independent replicates, values are means ± SEM. Two-sided Student’s t-tests were used for the two conditions. *P* Values at each time are indicated above the graph. Source numerical data and unprocessed blots are available in Source data.

In wild type ESCs, approximately one-third of Dppa2 binding sites are in H3K9me3 domains that predominantly mark L1/LINE elements and are enriched for the histone variant H2A.Z and open chromatin (**Fig. 8e and Extended Data** Fig. 10b). Annotating the published data, we found that Dppa2/4 deletion causes a loss of H2A.Z and chromatin accessibility specifically at H3K9me3 domains (**Fig. 8e)**. Upon degradation of H3K9me3 KMTs, Dppa2 footprinting, chromatin accessibility, and transcription of the associated transposable elements, especially LINE1 elements in H3K9me3 domains, are increased (**Fig. 8f,g and Extended Data** Fig. 10c). The increased Dppa2 binding in heterochromatin, upon dual degron treatment, correlates with fewer Dppa2 footprints and lower transcription at alternate developmental genes (**Fig. 8g**). We confirmed increased Dppa2 binding at several footprinted sites in the L1Md_T subfamily of LINE1 DNA repeats (**Extended Data** Fig. 10d) and knocking down Dppa2 dampened the activation of these DNA repeats (**Fig. 8h-i**). The findings highlight the critical role of H3K9me3- heterochromatin in balancing the genetic networks and DNA repeats that maintain pluripotency (**Extended Data** Fig. 10e).

## Discussion

Prior genetic deletions of the H3K9me3 KMTs in mammalian cell lines and mice required days for the deletions to take effect, making it difficult to assess how stable H3K9me3 domains are maintained across the genome at high temporal resolution^4,25,26^. Using acute degradation systems, we unveiled a dynamic nature of H3K9me3 maintenance in mammalian cells, “binary switches” controlling H3K9me3 heterochromatin integrity, and H3K9me3-dependent proteins and gene networks sensitive to H3K9me3 perturbations. In contrast to the prior single-locus studies, we discovered four classes of H3K9me3 stability types (**Fig. 3a**), in which different ChromHMM states with distinct proteins, histone variants/modifications, and transcription factor binding patterns predict each H3K9me3 stability type (**Fig. 3f-h**).

Notably, heterochromatin state 4, associated with highest H3K9me3 decay rate, is enriched for histone variants H3.3 and H2A.Z, transcription factor Dppa2, and the H3K9me3 demethylase KDM4C, and marks developmental genes and young L1Md_T (LINE1) subfamilies important for early development^49,50^. In contrast, heterochromatin state 1, associated with a lower H3K9me3 decay rate, is enriched for proteins that nucleate H3K9me3 heterochromatin at endogenous retroviral elements (ERV), including ZFP809^51^, TRIM28^52^, the m6A writer METTL3 and reader YTHDC1^36,37^. Furthermore, we note how certain heterochromatin proteins enriched in heterochromatin state 4 dissociate from chromatin faster than proteins enriched in heterochromatin state 1, suggesting varying dependencies of different heterochromatin- associated proteins on H3K9me3. The differential sensitivity to H3K9me3 erosion could lead to asymmetries in H3K9me3 inheritance between leading and lagging strand at different heterochromatin regions during DNA replication, as reported recently^53^, underscoring different stabilities in heterochromatin inheritance. Therefore, our study reveals how complex chromatin environments modulate H3K9me3 maintenance at developmental genes and repeat families and offers insights into how heterochromatin at developmental genes can be extensively remodeled, while maintaining repression of transposable elements and genome stabilities. The insights could be leveraged to modulate cell differentiation states at will with pre-determined impacts on repeat elements^54^.

Within 12 hours of KMTs’ loss, the rapid dissociation of HP1 proteins from H3K9me3 heterochromatin and a partial, but critical loss H3K9me3 lead to de-repression of diverse lineage programs and transposable elements, culminating in irreversible loss of pluripotency. Intriguingly, the chromodomain in HP1 and similar domains in other epigenetic readers, including the BAH domain, Tandem Tudor domain (TTD), Ankyrin repeats (in G9a/GLP), and EED WD40-repeats all have relatively low affinities towards their respective heterochromatin marks^9,55–58^. The low affinities of epigenetic readers allow for “binary switches”^59^, when neighboring residues are phosphorylated or when heterochromatin marks are reduced to a threshold level, as shown here, allowing transcription factors to quickly access partially de-compacted heterochromatin and leading to transcriptional de-repression, loss of the heterochromatin and exit from pluripotency (**Fig. 1i-j**). Concordantly, during mouse embryonic development, from day 4.5 to 6.5, SETDB1 and H3K9me3 are transiently reduced and cell proliferation is drastically accelerated^60^. Based on our findings, a transient reduction of HP1 and H3K9me3 is sufficient to reduce heterochromatin compaction, enabling pioneer transcription factor binding to create competence for multi-lineage specification in development.

Furthermore, the low affinity of heterochromatin reader proteins for epigenetic marks necessitates additional recruitment mechanisms. These include KMT dependencies and potential recruitment by heterochromatin protein TRIM28/KAP1, which may be directed by ZFPs at residual HP1β peaks, apparently independent of H3K9me3 marks. The design is reminiscent of the suboptimal TF motifs that increase the specificities of tightly controlled developmental enhancers^61^. Future analysis will test the functional significance of the underlying DNA sequences at the residual HP1 peaks in directing the establishment of H3K9me3 domains.

## Acknowledgments

We thank A. Barral and A. Katznelson for comments on the manuscript, and A. Hsieh, J. Lerner, R. McCarthy, and K. Kaeding for the valuable feedback during the project. J. Z. was supported by a Human Frontiers Science Program fellowship (LT000761/2019-L), and the research was supported by NIH R01GM36477 and R35GM153180 to K. Z.

## Author Contributions Statement

J. Z. and K. Z. conceived and designed the experiments and wrote the manuscript. J. Z. carried out the experiments and collected and analyzed the data. J. Z. and D. N. established the cTKO mouse ESC line, M.B.G carried out mass spectrometry sample preparations and data analysis. G. D. provided bioinformatic analysis, and T. L. and G. D. performed the mathematical modeling.

## Methods

### Mouse ESC derivation, culture, and maintenance

For this study, animals’ care was conducted in accordance with institutional guidelines. The derivation of mouse ESCs from blastocysts has been described previously^62^. Briefly, cTKO blastocysts were isolated at E3.5, and individual blastocyst was manually transferred onto irradiated MEF feeder cells in 4-well coated with 0.1% gelatin prepared one day before. The cells were cultured in 37 °C incubator with 5% CO2. The outgrowth of the ICM were expanded in 2i/LIF/FCS medium (DMEM-high glucose, 15% FBS, 2 mM GlutaMAX, 1 mM sodium pyruvate, 0.1 mM MEM NEAA, 0.1 mM 2-mercaptoethanol, 1,000 IU ESGRO, 1 μM PD0325901 and 3 μM CHIR99021) for 10 days before being dissociated with TryLE (Thermo, 12605010) and passaged 1 in 4 in 2i LIF/FCS medium for an additional 10 passages. From passage 11, ESCs were cultured in LIF/FCS culture conditions (without PD0325901 and CHIR99021) in feeder-free conditions. To passage the cells, mESCs were dissociated with TryPLE at 37 °C for 2 minutes, and resuspended in 4x volumes of mESC medium, and pelleted by centrifugation at 200x g for 3 minutes. Cell pellets were resuspended in 4 ml mESC medium, counted, and plated at 24,000 - 40,000 cells/cm^2^. All ESC lines were routinely tested for mycoplasma contaminations.

### Generation of cTKO and dTKO mouse ESC line

The established cTKO mouse ESCs were used for generating the cTKO-creERT2 and dTKO degradable mouse ESC line. The plasmid with CAG promoter driving creERT2-IRES-Bsr construct (addgene, #48760) was linearized, ethanol precipitated, resuspended in nuclease-free H2O, and transfected into mouse cTKO ESCs with lipofectamine 3000 at 2.5:1 ratio (2.5ul of lipofectamine 3000 and 1 μg of linearized plasmid). The transfected cells were allowed to recover for 2 days, before being re-plated at clonal density in the mESC medium supplemented with 10 μg/ml Blasticidin S hydrochloride (Thermo, R21001) for 7 days. The blasticidin resistant colonies were picked and transferred to 48-wells and expanded further.

The knock-in targeting vectors were constructed using a strategy described previously^63^. The targeting vector for inserting atAFB2 to the *TIGRE* safe harbor was modified from the plasmid pEN396, (addgene, #92142) by replacing the Tir1 inserts with the atAFB2-NLS sequence^27^ amplified from addgene plasmid #129717. The targeting vector for inserting mini-IAA7 to the *Suv39h1* allele was constructed using Gibson assembly (NEB, E2611), with addgene plasmid #86233 as the backbone. The 1 kb Suv39h1 homologies were amplified from the mouse genomic DNA, mini-IAA7 was amplified from addgene plasmid #129721, the FRT flanked selection cassette FRT-PGK-neo-FRT was amplified from addgene plasmid #86233. The targeting vector for inserting dTAG-Setdb1 full-length CDS to the *Setdb1* alleles was constructed using Gibson assembly. The 1 kb Setdb1 homologies and Setdb1 3’UTR were cloned from mouse genomic DNA, the bsr selection gene was amplified from addgene plasmid #92140, dTAG domain^28^ was amplified from addgene plasmid #91798, and Setdb1 CDS was synthesized by GeneScript. The 3 tandem polyA sequences^29^ were amplified from addgene plasmid #61576. The gRNAs were cloned into eSpCas9(1.1) (addgene, #71814) using BbsI digestion. The list of gRNA sequences used in this study is included the in supplementary materials.

For knocking-in experiments, the Cas9 plasmid co-expressing gRNAs and targeting vectors were co-transfected with lipofectamine 3000 into mouse ESCs cultured in 6-well plate. Cells medium were replenished daily for 2 days, before being dissociated into single cells with TryLE and plated at 10,000 cells/10cm dish and selected in the next day for 7 days with appropriate drugs until the drug-resistant colonies emerges. Normally, 50 colonies were manually picked in each round of targeting, and each clone genotyped while expanding them.

For the degradation timecourse, mouse ESCs were plated at 67,000 cells/cm^2^ and allowed to attach overnight. The auxin and dTAG13 were added to the medium at the final concentration of 100 μg/ml and 0.5 μM, respectively for specific durations before the cells were collected for downstream analysis. The degrons were replenished daily for time course experiments that require more than 24 hr of treatment.

### Thymine treatment and KDM4 inhibition or knockdown

For blocking DNA synthesis, one million cells were plated in a 6-well one day before, and 1mM thymidine was added to the medium 6 hours before the degradation time course and was replenished every 24 hr. The complete block of DNA synthesis was verified with EdU integration. Cells were incubated with 20 µM EdU added to the medium 2 hr before being dissociated and fixed with 4% PFA. The fixed cells were then permeabilized in 0.3% Triton in DPBS for 10 mins, followed by click-it chemistry reactions using Click-iT™ EdU Cell Proliferation Kit (Thermo, C10337). The EdU integration was analyzed with BD Accuri C6 Flow Cytometer. For KDM4 inhibition, 100 nM QC6352 (MedChemExpress, HY-104048) was added to the medium 2 days before the degron treatment and was replenished every 24 hr.

For the KDM4A-C knockdown, mouse ESCs were transfected with SMARTpool siRNAs for KDM4A (Dharmacon, 230674), KDM4B (Dharmacon, 193796), and KDM4C (Dharmacon, 76804) at 6.7 nM final concentration for each siRNA or non-targeting control siRNA (Thermo, Silencer Select, 4390843) at 20 nM final concentration using Lipofectamine RNAiMAX Transfection Reagent (Thermo, 13778150) following the manufacturer’s instructions. The cells were allowed to recover for 2 days before treating with auxin and dTAG degrons for different durations.

### Protein analysis

To ensure the accuracy of the timings of degron treatment, cells were directly lysed in Trizol reagents (Thermo, 15596026), and protein was purified following manufacturer’s instructions. For less time-sensitive protein analysis, cells were directly lysed in RIPA buffer (50 mM Tris, 150 mM NaCl, 1% NP-40, 0.5% of Sodium Deoxycholate, 0.1% SDS) supplemented with 1x complete proteinase inhibitor. After 30 minutes of incubations in RIPA buffer on ice, the whole cell lysates were sonicated with Bioruptor at high power for 10 minutes with 30 seconds on and 30s off intervals, followed by centrifugation for 20,000x g at 4 °C for 20 minutes. The soluble proteins were then transferred to a new tube and quantified with BCA method and stored in -80 °C for long-term storage.

For western blot analysis, cell lysates were mixed with 1x LDS Sample Buffer (Thermo, NP0008) and 1x Sample Reducing Agent (Thermo, NP004) and boiled for 5 minutes. Ten μg of protein per well were loaded and transferred to PVDF membrane, followed by 1 hour incubation with blocking buffer (5-10% non-fat milk in TBS) at room temperature and over-night incubations with primary antibody diluted in blocking buffer with 0.1 % Triton at 4 °C. The membrane was washed 3x in wash buffer (TBS with 0.1% Triton X-100), followed by 1 hour incubation with secondary antibodies at room temperature and 3x washes, and visualized on Amersham Imager 680. The antibodies and their dilutions used in this study are as follows: H3K9me3 (Abcam, ab8898, 1:10,000 dilution); H3 (Abcam, ab1791, 1:20,000); H4K20me3 (Millipore, 07-463, 1:2000); H3K9me2 (Abcam, ab1120, 1:5000); H3K27me3 (Millipore, 07-449, 1:5000); NF-YA (Santa Cruz, sc-17753. 1:1000); α-Tubulin (Cell Signaling Technology, 2144, 1:5000); Dppa2 (Millipore, MAB4356, 1:2000); Kdm4a (Thermo Fisher, PA5-14782, 1:2000); Kdm4b (Bethyl, A301-478A, 1:2000); Kdm4c (Gifts from the Kristian Helin Lab, 1:2000); HP1α (Cell Signaling Technology, 2616, 1:5000); HP1β (Cell Signaling Technology, 8676, 1:5000); HP1γ (Cell Signaling Technology, 2619, 1:5000); Mpp8 (Proteintech,16796-1-AP, 1:2000); TRIM28 (Abcam, ab22553, 1:1000); YTHDC1 (Cell Signaling Technology, 77422, 1:2000); Mettl3 (Bethyl, A301- 567A, 1:2000); HA (BioLegend, 901533, 1:2000), V5 (eBioscience, 14-6796-82, 1:2000), Suv39h1 (Thermo Fisher, 702443, 1:1000), Setdb1 (Proteintech, 11231-1-AP, 1:1000); Donkey anti-mouse-HRP (Jackson InnumoResearch laboratories, 715-035-150, 1:5000); Donkey anti-Rabbit-HRP (Jackson InnumoResearch laboratories, 711-035-152, 1:5000); Donkey anti-goat-HRP (Jackson InnumoResearch laboratories, 705-035-003, 1:5000).

### Chromatin Fractionation

The chromatin fractionation was performed as previously described with minor modifications^64^. Briefly, 20 million mouse ESC cells were used as starting material and each sample was prepared in triplicates. Cell pellets are resuspended in swelling buffer (10 mM HEPES pH 7.9, 1.5 mM MgCl2, 10 mM NaCl, 1 mM DTT, 0.15% NP-40) freshly supplemented with 1 mM DTT and 1x PIC and incubated for 10 minutes on ice. The nuclei were pelleted by centrifugation for 10 min at 800x g at 4 °C, and cytoplasmic extracts were collected in separate tubes. The nuclei pellets were washed once with swelling buffer, and resuspended in nuclei lysis buffer (10 mM Tris-HCl (pH 7.0), 4 mM EDTA, 0.3 M NaCl, 1 M urea, and 1% (vol/vol) NP-40) with 1mM DTT and 1x PIC for 15 minutes on ice, and gently vortexed briefly, and chromatin fraction were pelleted by centrifugation for 5 minutes at 1000x g at 4 °C, and nucleoplasmic extracts were collected in separate tubes. The chromatin pellets were washed twice with nuclei lysis buffer without EDTA, and resuspended in MNase digestion buffer (20mM HEPES pH7.5, 500mM NaCl, 3mM CaCl2, 0.3% NP-40) freshly supplemented with 0.5mM DTT, 1x PIC and 100 units of MNase per sample and incubated at 37 °C for 10 minutes with rigorous shaking. The MNase digestion were stopped by adding EDTA to the reaction at final concentration of 10mM and solubilized chromatin were collected by centrifugation at 18000 xg for 20 minutes at 4 °C.

### Mass spectrometry analysis of histones and chromatin fractions

All mass spectrometry samples were collected in triplicates at each time-point. The dTKO mouse ESCs were treated with auxin and dTAG for different durations, and cells were collected by scrapping in cold room, and pelleted by centrifugation at 4 °C, 500x g for 5 minutes, and snap frozen in -80 °C for storage.

Histone purification was performed using 5 million mouse ES cells following the protocol as previously described^65^. Briefly, cell pellets were thawed and incubated in nuclei isolation buffer (15mM Tris–HCl (pH 7.5), 15mM NaCl, 60mM KCl, 5mM MgCl2, 1mM CaCl2, 250mM sucrose, 0.3% NP-40) freshly supplied with 1mM DTT, 0.5mM AEBSF and 10mM sodium butyrate, and nuclei were separated by centrifugation (600 g for 5 min), followed by 50 µL of cold 0.4N H_2_SO_4_ treatment at 4°C with shaking for 3 hours without rotation and pelleted at 14000 g for 10 min. Histones were then precipitated by adding TCA to a final concentration of 33% (w/v), and purified histones were resuspended in 20ul of 50mM NH_4_HCO_3_, pH 8.0, followed by derivitization and digestion using the bottom-up strategy described previously (Sidoli et al., 2016). Briefly, 15ul of pure propionic anhydride:acetonitrile (1:3) mix was added to the histone samples to allow derivatization of the lysine residues side chains, followed by addition of 3ul of ammonium hydroxide to adjust the pH to 8.0 immediately after adding propionic anhydride to the mixture. The reaction was repeated once. Protein was digested with trypsin overnight at an enzyme:sample ratio of 1:20 at room temperature. The derivatization reaction was repeated to derivatize peptide N-termini, followed by desalt using Pierce C18 Tips, 10 µL bed (Thermo Scientific).

Both histone tail and chromatin-fractions peptide peptides were analyzed with nanoLC-MS/MS, as previous described^65^. For histone analysis, peptides were separated using an UltiMate3000 (Dionex) HPLC system (Thermo Fisher Scientific, San Jose, Ca, USA) using a 75 μm ID fused capillary pulled in-house and packed with 2.4 μm ReproSil-Pur C18 beads to 20 cm. For histone analysis, the HPLC gradient was set at a flow rate of 300nl/min with 0%–34% solvent B (A = 0.1% formic acid, B = 95% acetonitrile, 0.1% formic acid) over 46 minutes and from 34% to 90% solvent B in 5 minutes. The QExactive HF (Thermo Fisher Scientific, San Jose, CA, USA) mass spectrometer was configured using the data-independent acquisition (DIA) method. Full scan MS spectra (m/z 300-1100) were acquired with a resolution of 60,000 with an AGC target of 1e6; MS/MS spectra were obtained with 50 m/z precursor isolation windows (stepped CID normalized collision energy of 25, 27.5, 30 and an AGC target of 5e5. Mass spectrometry data were imported into Skyline^66^ to calculate integrated MS2 peak areas. The data was median normalized and for statistical analysis, a 2-tailed student t-test was performed (significant if p < 0.05).

For chromatin-fraction, the HPLC was set at a flow rate of 300nl/min with gradient 0%– 35% solvent B (A = 0.1% formic acid, B = 95% acetonitrile, 0.1% formic acid) over 90 min and from 45% to 95% solvent B in 30 min. A data-independent acquisition (DIA) chromatogram library method was used to configure the QExactive HF (Thermo Fisher Scientific, San Jose, CA, USA) mass spectrometer. Full scan MS spectra (m/z 385-1015) were acquired with a resolution of 60,000, AGC target of 1e6 for wide-window data and 8 m/z staggered isolation windows (normalized collision energy of 27.5 and an AGC target of 1e6) were used to acquire MS/MS spectra. The chromatogram library was collected in 6 gas-phase fractions (GPF) per fractionated compartment with DIA with full scan MS spectra of 110 m/z each with 4 m/z overlapping windows with the same resolution and AGC targets settings as the wide window data^67^. Chromatogram library was built from 6 gas-phase fractions using Walnut in EncyclopeDIA. Mass spectrometry data files were demultiplexed with MSConvert^68^ and wide-window data was processed in EncyclopeDIA^67^ with chromatogram library from Walnut. Processed mass spectrometry data files were imported into Skyline^66^ and, which was then exported to MSstats^69^ and the raw count data was analyzed with RUVseq package^70^. Briefly, to increase the robustness of the analysis, proteins that have more than 4 NAs (undetected) across the time series suggest unreliable detections and therefore were removed from the table of raw counts. Next, to determine negative controls that remain constant during the time course, using RUVg function we first empirically determined top 20 proteins that shows the least variations across the time series, including H3, H4, Hnrnph1, and Polr2c. These protein lists were then used as empirical internal controls for RUVseq normalization function RUVg (empirical, k =1). The normalized protein signals were then used for downstream analysis and visualization.

### Chromatin Immunoprecipitation

Chromatin immunoprecipitation was performed as previously described^71^. Briefly, cells were crosslinked with 1% formaldehyde at room temperature for 10 mins and quenched with 0.125 mM Glycine at room temperature for 5 minutes. Cells were then washed 3 times with ice cold DPBS and incubated with 7 ml swelling buffer (25mM HEPES-KCl, pH7.2, 1.5 mM MgCl2, 20mM KCl, 0.1% NP-40) per 15 cm dish for 10 minutes at 4 °C. The cells were scrapped off the plates, collected and pelleted in 15 ml falcon tubes and snap frozen in the liquid nitrogen, and stored in - 80 °C freezer.

On the day of sonication, the crosslinked cells were incubated with lysis buffer 1 (50 mM HEPES-KOH, pH7.5, 140 mM NaCl, 1mM EDTA, 10% glycerol, 0.5% NP-40, 0.25% Triton X-100, 1 mM DTT, and 1x complete protease inhibitors cocktail) and lysis buffer 2 (10 mM Tis-HCl, pH8, 200 mM NaCl, 1mM EDTA, 0.5 mM EGTA, 1x complete protease inhibitors cocktail) for 5 minutes each at 4 °C. The nuclei were then resuspended in the lysis buffers 3 (10 mM Tis-HCl, pH8, 200 mM NaCl, 1mM EDTA, 0.5 mM EGTA, 1 mM DTT, 0.1% Na-Deoxycholate, 0.5% N- lauroylsarcosine, 1x complete protease inhibitors cocktail) for 10 minutes on ice, followed by centrifugation at 1350g for 5 minutes. The nuclei pellets were then resuspended in lysis buffer 3 and sonicated with Covaris (Incident peak power: 140; Duty Cycle: 5; Burst/Cycle: 200) for 15 minutes. The sonicated chromatin was centrifuging at 20,000g for 20 minutes and the soluble chromatin were collected, and 20 μl chromatin was de-crosslinked in 200 μl elution buffer (50 mM Tris-HCl, pH 8.0, 10 mM EDTA, 1% SDS) at 65 °C over night and the rest of the chromatin was aliquoted, snap frozen in liquid nitrogen and stored in -80 °C freezer. The de-crosslinked chromatin was treated with proteinase K and RNase, followed by ethanol precipitation and DNA was resuspended in nuclease-free H2O. For H3K9me3 and HP1β ChIP-seq, 20 μg of chromatin was combined with 12.5 ng of Drosophila spike-in chromatin (0.06% of mouse chromatin) and incubated with 10 μl of Protein A/G dynabeads pre-coated with 0.5 μg of H3K9me3 (abcam, ab8898) or 1ug of HP1β antibody (Cell Signaling Technology, 8676) and 0.5 μg of drosophila spike-in antibody on a rotating wheel at 4 °C overnight. For H4K20me3 (Abcam, ab8896), H3K9me2 (Abcam, ab1120), H3K27me3 (Millipore, 07-449), TRIM28 (Abcam, ab22553), YTHDC1 (Cell Signaling Technology, 77422), and Mettl3 (Bethyl, A301-567A), Dppa2/4 (R&D

Systems, AF3730), 1 μg antibody were used per 20 μg of mouse chromatin. The chromatin-antibody-dynabead slurry was then washed 5 times with RIPA buffer (50 mM Hepes-KOH, pH 7.6, 500 mM LiCl, 1mM EDTA, 1% NP40, 0.7% Na-Deoxycholate, 1x complete protease inhibitors cocktail) and once with TE (10 mM Tris-HCl, pH 8.0, 1mM EDTA). The chromatin was eluted in elution buffer by incubating at 65 °C for 20 minutes and de-crosslinked at 65 °C overnight, followed by proteinase and RNase treatment and ethanol precipitation. The DNA concentration was quantified with Qubit HS DNA kit and stored in -20 °C.

ChIP-seq DNA libraries were prepared with NEBNext® Ultra™ II DNA Library Prep Kit (NEB, E7645) following the manufacturer’s instructions. Briefly, 10 ng of ChIP DNA was used for each library and amplified with 11 PCR cycles and the library DNA was purified with Ampure XP beads and the distribution of the libraries DNA fragments were analyzed using High Sensitivity D1000 DNA ScreenTape (Agilent, 5067-5584) on Agilent Tapestation 4150.

### ChIP-seq data Analysis

The mouse mm10 and Drosophila dm6 genome were merged to generate a hybrid genome, and the bowtie2 index for the hybrid genome was generated using ‘bowtie2-build -f’. The ChIP-seq fastq files were aligned to the hybrid genome with: ‘bowtie2 -p 16 --local -X 1000 -S output.sam’ and the unmapped and unpaired reads were filtered out using ‘samtools view -bS -h -f 3 -F 4 -F 8’. Since the H3K9me3 heterochromatin and HP1 are highly enriched in repetitive DNA, the multimappers were included in the analysis. The drosophila spike-in normalization strategy controls for the chromatin immunoprecipitation and library amplification steps, therefore, the percentage of Drosophila reads in each ChIP sample were used to calculate the scalars for normalization. The bigwig files were generated using deeptools bamCompare with log2 ratio of ChIP/Input and scaled based on the ratio of the spike-in reads. For H3K9me3 domain calling, the bam files of the triplicates of each sample were merged, which were then converted into bed files using bedtools bamtoBed. H3K9me3 domains were called using RSEG^35^ ‘rseg-diff -c mm10.chrom.sizes.bed -o RSEG.bed -i 20 -v -mode 2 -d deadzones-k36-mm10.bed chip.bed input.bed’. ChIP signals in each domain were quantified using the bigWigAverageOverBed and the mean values were used for downstream analysis. To generate alluvial clusters, H3K9me3 levels at 43,000 randomly sampled 3 kb windows (median size of H3K9me3 domains) outside H3K9me3 domains were quantified using bigWigAverageOverBed to calculate the background H3K9me3 levels in the genome. The inflection point (at 0.35) of H3K9me3 signal distributions between RSEG H3K9me3 domains and randomly sampled 3 kb windows were used as the cut- off to determine the timing of the H3K9me3 loss over the degradation time course.

### ATAC-seq

ATAC-seq were performed after the ChIP-seq time-series. Since ChIP-seq analysis shows little H3K9me3 changes within 1 hr, the 1hr time-point is omitted from sample preparations for the ATAC-seq analysis. Instead, 6 hr time-point is added to analyze the transcription factor footprinting at finer details. The ATAC-seq was performed in triplicates following the protocol described previously^72, 73^ with slight modification. Briefly, 100,000 cells were collected and washed once with DPBS, and nuclei were isolated by incubating the cells with 50 μl ice-cold ATAC-RSB buffer (10 mM Tris-HCl pH 7.6, 10mM NaCl, 3mM MgCl2) with 0.1% NP40, 0.1% Tween 20, and 0.01% Digitonin for 3 minutes on ice. The nuclei were then washed once with ATAC-RSB buffer with 0.1% Tween 20 and incubated with 50 μl transposition mix (25 μl 2x TD buffer, 2.5 μl transposase (120 nM final), 16.5 μl PBS, 0.5 μl 1% digitonin, 0.5 μl 10% Tween-20, 5 μl H2O) at 37°C for 30 minutes and mixed in thermomixer at 1000 RPM. The transposition was stopped by adding 250 μl (five volumes) DNA binding buffer to the transposition mix and DNA was purified with Zymo DNA Clean and Concentrator-5 Kit (D4014) and eluted in 21 μl elution buffer. For library preparation, 5 cycle of pre-amplification was performed, and the DNA concentration was then quantified, followed by an additional 5 cycles of amplifications. The library DNA was purified with Zymo DNA Clean and Concentrator-5 Kit and eluted in 20 μl of H2O.

ATAC-seq data was aligned to mouse mm10 genome, and the analysis was performed with CUT-RUNTools-2.0 ^74^ (https://github.com/fl-yu/CUT-RUNTools-2.0) using the standard CUT&Tag pipelines with “120bp fragment” OFF. Briefly, the FASTQ files were trimmed with “ trimmomatic PE -phred33 ILLUMINACLIP: Truseq3.PE.fa:2:15:4:4:true LEADING:20 TRAILING:20 SLIDINGWINDOW:4:15 MINLEN:25”, and then aligned to mouse mm10 genome with “bowtie2 --very-sensitive-local --phred33 -I 10 -X 700”, Unmapped reads were filtered out using “samtools view -bh -f 3 -F 4 -F 8”. Considering that significant amounts of regions that gained ATAC signals are within the repeats, including many transposable elements, we kept multimapping reads for the downstream analysis. The narrow peaks were called for each sample, and all the peaks were then merged and used for downstream analysis. When generating the bigwig files, the reads were first shifted using “”alignmentSieve –ATACshift --bam” that shifts the plus strand reads by +4 bp, and minus strand reads by -5 bp, before using deeptools bamCoverage with “CPM” normalization to generate bigwig file.

The motifs of all vertebrate transcription factors were downloaded from the JASPAR database (https://jaspar.genereg.net/) and the ATAC footprints of all the transcription factor motifs were performed using TOBIAS with the default settings (https://github.com/loosolab/TOBIAS).

Then the motifs of the transcription factors that are expressed in mouse ESCs were selected based on our TTseq data: only the transcription factors that has at least 10 reads mapped to the gene bodies at any time point were used for downstream analysis. The nucleosome positioning analysis from the ATAC data was performed using NucleoATAC (https://github.com/GreenleafLab/NucleoATAC) with default settings. The bedgraphs from the smoothed nucleoatac signals were converted into bigwig using bedGraphToBigWig. The resulting bigwig files were used to generate the heatmaps of nucleosome positions surrounding the transcription factor footprinted sites using deeptools.

### Nascent RNA extraction and TT-seq

TT-seq experiments were performed after the H3K9me3 ChIP-seq time course. ChIP-seq time series shows little H3K9me3 changes within 1 hr, therefore the 1hr time-point is omitted from sample preparations for the TT-seq. TT-seq was performed in triplicates as previously described^75, 76^ with slight modifications. Cells were labeled with 4SU (Glentham Life Sciences, GN6085) at 1mM final concentration for 10 minutes in 37 °C incubator. The medium was then immediately decanted, and cells were lysed by directly adding 5 ml Trizol reagent onto the plates. Cells were scrapped from the plates and collected in 15 mL falcon tubes and frozen in -80 °C. Total RNA were extracted using Trizol RNA extraction protocol following the manufacturer’s instructions and stored in -80 °C. S. cerevisiae 4-thiouracil (4TU, Sigma, 440736) labelled RNAs were prepared following the protocol described previously^76^.

For TT-seq, 110 μg total RNA, combined with 1 μg of spike-in S. cerevisiae 4TU-labeled RNA were fragmented with 166 mM NaOH for 30 minutes on ice, and stopped by adding 400 mM Tris-HCl, pH 6.8 and passed through Micro Bio-Spin P-30 gel columns twice. The total RNAs were then denatured at 65°C for 10min, followed by 5mins of incubation on ice, and mixed with 50 μL of 0.1 mg/ml MTSEA biotin-XX linker (Biotium, BT90066) on rotator at room temperature for 30 minutes in the dark. The RNA was then purified with phenol/chloroform/isoamyl alcohol (25:24:1 (vol/vol/vol)), followed by isopropanol precipitation. The RNA was then resuspended in nuclease-free H2O and denatured at 65°C for 10min, followed by 5 minutes of incubation on ice and mixed with 150 μL of μMACS streptavidin MicroBeads (Miltenyi, 130-074-101) for 15 minutes in cold room on a rotating wheel. The biotinylated RNA was retained on μMACS magnetic separator (Miltenyi, 130-042-602) placed on a MACS multistand (Miltenyi, 130-042-303) and washed three times with 900 μL of pre-warmed (65 °C) pull-down wash buffer (100 mM Tris-HCl, pH 7.4, 10 mM EDTA, 1 M NaCl and 0.1% Tween 20) and three more times with 900 μL of room temperature pull-down wash buffer. The biotinylated RNA was eluted twice from the column with 100 μl of 100 mM DTT and purified with RNeasy MinElute Cleanup Kit (Qiagen, 74204).

TT-seq libraries was prepared with NEBNext® Ultra™ II Directional RNA Library Prep Kit (NEB, E7760) following the manufacturer’s instructions. 21 ng of nascent RNA was used for each sample and amplified with 11 PCR cycles.

### TT-seq analysis

For TT-seq analysis, the mouse and yeast hybrid genome were generated by merging the mouse mm10 and sarCer3 fasta files and the hybrid genome index were built with STAR (2.5.2a version) using “STAR --runMode genomeGenerate --genomeDir index_director –genomeFastaFiles mm10_sarCer3_hybrid.fa --sjdbGTFfile Hybrid_mm10_sacCer3_hybrid.gtf --sjdbOverhang 41”. The TT-seq fastq files were aligned to the hybrid genome using the STAR alignment parameters recommended by TEtranscripts analysis pipeline^77^ (https://github.com/mhammell-laboratory/TEtranscripts) “--winAnchorMultimapNmax 100 --outFilterMultimapNmax 100”. The percentage of the reads aligned to the yeast genome in each sample were used to calculate the scalar for bigwig and DEseq2 normalization between different samples. The transcription units were called using TU_filter (https://github.com/shaorray/TU_filter). The expression level of the genes, non-coding RNAs and transposable elements were quantified with TElocal (https://github.com/mhammell-laboratory/TElocal) with “TElocal --TE mm10_rmsk_TE.gtf.locInd - -stranded reverse” and analyzed with DEseq2 R package with spike-in scalar calculated above. The bigwig files were generated with deeptools bamCoverage using RPKM normalization, with forward and reverse strands separated.

To integrate the H3K9me3 and HP1β ChIP-seq data, ATAC-seq and TT-seq data, first the H3K9me3 domains and ATAC peaks were intersected using bedtools, with at least one bp overlap. Then the nearest ATAC peaks to the transcription units (TUs) in the TT-seq were defined using bedtools closestBed. Only the ATAC peaks-TU pairs that are less than 10 kb in distance were considered as being associated and used for downstream analysis, including supervised heatmaps. The supervised heatmaps were generated with pheatmap R package.

### High-throughput sequencing

All NGS libraries were sequenced in paired end with 42 reads at each end using NSQ 500/550 Hi Output KT v2.5 (75 CYS) kit, on Illumina NextSeq 500.

### ChromHMM modeling

The ChromHMM model was generated using the procedure published previously^78^ with default settings. The fastq files were downloaded from GEO and aligned to mouse mm10 genome using parameters described above. The following ChIP-seq datasets were used to construct the ChromHMM model: H3K79me2 (GSM2417104), H3K36me3 (GSM2417108), H3K4me1 (GSM2417088), H3K4me2 (GSM2417084), H3K4me3 (GSM2143325), H3K27me3 (GSM2417100), H2AK119ub1 (GSM3379157, GSM3379158, GSM3379159), H3K27Ac (GSM4830129, GSM4830130, GSM4830131), H3K122ac (GSM3143871), H3K64ac (GSM3143869), ATAC (GSM2417076), Ctcf (GSM2609185, GSM2609188), RNA Pol2 (GSM4874306, GSM4874307), Med23 (GSM4874310, GSM4874311), H3.3 (GSM2080325, GSM2080330, GSM2080335), H4K20me3 (GSM4664503), ATRX (GSM1372574, GSM1372576), TRIM28 (GSM3517473, GSM3517474, GSM3517475), Mpp8 (GSM5393701, GSM5393704) Mttl3 (GSM3594077, GSM4664507), SETDB1 (GSM3594122, GSM4664499), YTHDC1 (GSM4664495, GSM4664569), ADNP (GSM2582357, GSM2582358, GSM2582359), Jmjd2c (GSM2460996, GSM2460997), SUMO2 (GSM2629947, GSM2629948), Chaf1a (GSM1819193), Brg1 (GSM2417177), ZFP809 GSM1819195), HDAC1 (GSM2417173), SMARCAD1 (ERR2699918), Suv39h1 (GSM1375157), Suv39h2 (GSM1375158), H3K9ac (GSM2417092), H3K9me3 (this study), H3K9me2 (this study), HP1β (this study).

### Mathematical modeling of H3K9me3 decay

The model was constructed using parameters described previously ^24^. The model is based on a discrete one-dimensional array of 1000 nucleosomes, where each nucleosome can have one of two states, unmodified (with a value of 0), or modified (with a value of 1). The model consists of three processes that govern the state of a nucleosome: nucleation and propagation, both of which are governed by the rate constant K+, and turnover, a process that can turn a nucleosome from a modified back to an unmodified state and governed by the rate constant K-. Each of the three processes proceeds by probabilistic description. Each process proceeds based its probability of occurrence *p*, given by the following equations for nucleation and propagation, *p*_*nuc*_ = *k*_+_Δ*t*, *p*_*prg*_ = *k*_+_Δ*t*, And for turnover, *p*_*trn*_ = *k*_−_Δ*t*. For simplicity, we chose to maintain Δ*t* = 1, such that the value of K+ and K- reflect the probability of the processes. In addition, consistent with a previous model ^24^, κ, the ratio between K+ and K- was introduced: 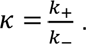

The simulation involves two steps: forward and reverse stage. In forward stage: H3K9me3 nucleates and propagates from an unmethylated state, and turnover was allowed to proceed until the number of modified nucleosomes achieves a steady state. Here we swept the values of K- between 0.0335 and 0.184, and κ between 1 and 1.6, the value of K+ was calculated accordingly. This stage was allowed to proceed for 15,000-time steps to ensure the model achieve the steady state. Reverse stage: After 15,000-time steps, the value of K+ was set to 0, allowing only turnover to proceed. The configuration of the array at the last time step of the forward stage served as the initial condition for the reverse stage. The lattice was allowed to return to a state where all nucleosomes were unmodified, for which 500 additional time steps were sufficient for all κ values. The reverse methylation simulation was fitted to an exponential decay curve with two characteristic degrees of freedom, *N* = *N*_0_ exp(−*a*Δ*t*), where *N*_0_ is the number of modified nucleosomes at the end of the forward stage (and therefore the beginning of the reverse stage), and *a* is the decay constant of the exponential curve. The simulation was repeated 2000 times for every k- and K pair, and curve parameters *N*_0_ and *a* were fitted for every simulation repeat.

To directly compare the experimental data with the results from the simulation, the H3K9me3 domains were broken into 200 bp bins, approximately the size of one nucleosome, and H3K9me3 signals within each 200 bp bin was quantified. Using the cut-off for background H3K9me3 levels mentioned above, each 200 bp bin is either “methylated” (set as 1), or unmethylated (set as 0). In this way, the experimental data was binarized to match the data format produced from the simulation.

The *N*_0_ and *a* parameters for every H3K9me3 alluvial cluster in the experimental data was compared to the simulated *N*_0_ and *a* for every pair of κ and K-. The best fit κ and K- pair was found using a combined least mean square objective function,

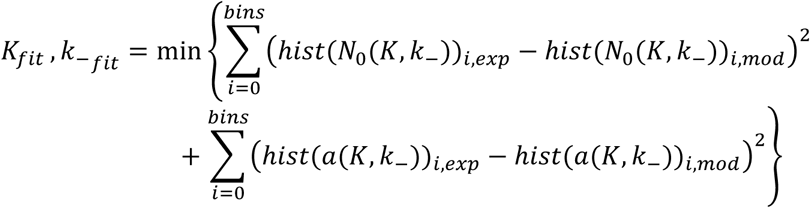

In the unique case of the late H3K9me3 cluster, two separate unique clusters were detected. The sub-clusters were first separated using an unsupervised Gaussian Mixture Model classifier, after which simulated *N*_0_ and *a* histograms were fitted to each cluster separately.

### Data availability

All the raw sequencing and processed data are available at the Gene Expression Omnibus under the accession number GSE233041. The mouse genome (mm10 assembly), the drosophila genome (dm6 assembly), and Yeast (S. cerevisiae) genome (sacCer3 assembly) are downloaded from UCSC genome browser: https://hgdownload.soe.ucsc.edu/downloads.html. The raw data supporting the findings of this study are included in the manuscript and the supplementary files. A list of publicly available datasets used in the study can be found in the manuscript.

### Code Availability

The parameters for next generation sequencing data analysis are reported in the method sections and included in GEO upload. All the original code for mathematical modeling of H3K9me3 decay has been deposited to the github: https://github.com/Tomer-Lapidot/H3K9me3_Methylation_Stochastic_Model.

### Statistics and Reproducibility

All experiments, including ChIP-seq, ATAC-seq, TT-seq, mass spectrometry, western blot and RT-qPCR were repeated three times independently, unless otherwise specified. Detailed statistical analysis were indicated in the figure legends and exact p values were reported. No statistical method was used to predetermine sample size. No data were excluded from the analyses. The experiments were not randomized.

## Supplementary Figure Legends

**Supplementary Fig. 1:**
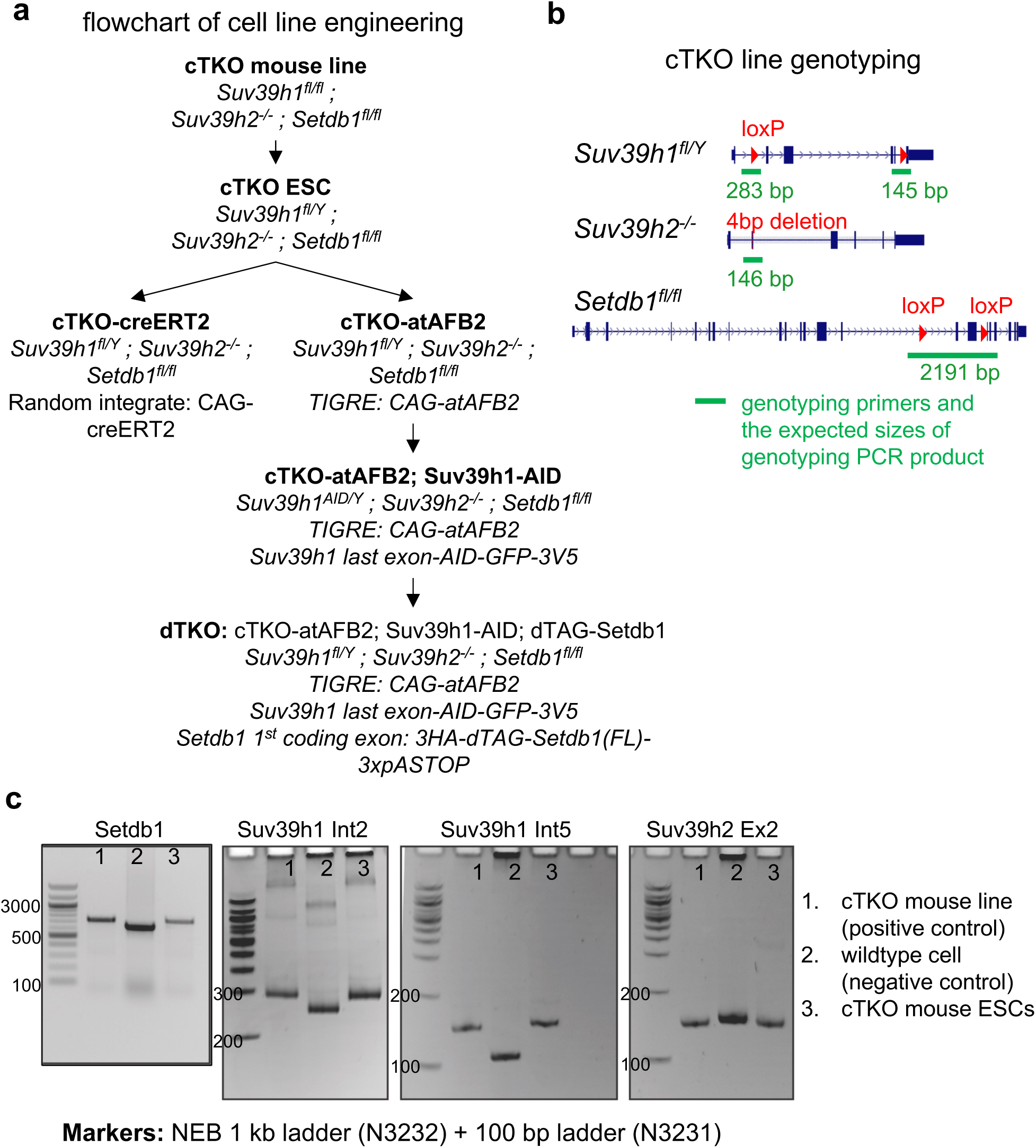
Establishing and genotyping cTKO mouse ESC line. **a**. Flowchart of the derivation, genetic engineering, and genotypes of cTKO and dTKO mouse ESC lines. cTKO, conditional triple knockout; dTKO, degradable triple knockout. **b.** cTKO mouse ESC genotypes, with the genotyping PCR primers targeted sequences and the expected size of PCR product highlighted in green. **c.** Genotyping of *Setdb1*, *Suv39h1* and *Suv39h2* cTKO mouse ESCs with primers indicated in **(b)**.

**Supplementary Fig. 2:**
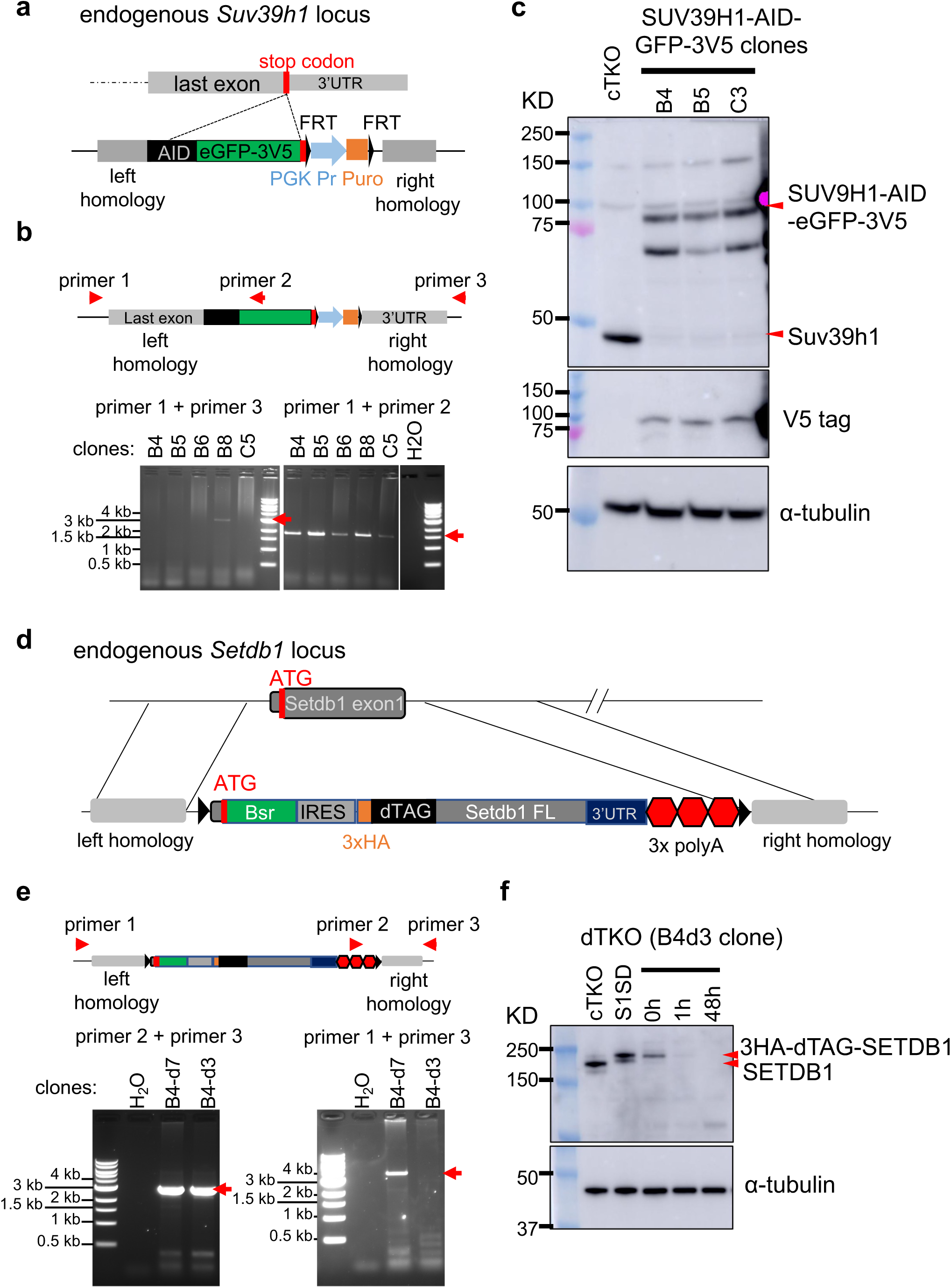
Genetic degron engineering of *Suv39h1* and *Setdb1* alleles. **a**. Schematics depicting the targeting strategy to insert AID-eGFP-3xV5 tags to the last exon of *Suv39h1* allele immediately before the Stop codons. AID, auxin inducible degradation; FRT, flippase recognition target; PGK Pr, PGK promoter; puro, Puromycin resistant gene. **b**. Top, schematics of *Suv39h1* genomic regions after degron engineering, with genotyping primers used for screening engineered clones indicated. Bottom, Genotyping of successfully targeted clones using primer pairs indicated at the top. The expected sizes are detailed in the supplementary table and was denoted with a red arrow. **c**. western blot of Suv39h1, V5 and loading control, αTubulin, of parental cTKO ESCs and Suv39h1 degron engineered clones, B4, B5 and C3. The molecular weight in kilodalton (KD) is indicated on the left, and the expected band of wildtype Suv39h1 and Suv39h1-AID-eGFP-3xV5 fusion proteins are indicated with red arrows on the right. **d**. Schematics depicting the targeting strategy to insert dTAG-Setdb1(full-length) to the first protein coding exon of *Setdb1* alleles. Bsr, blasticidin S-resistance gene; IRES, internal ribosomal entry site; FL, full length. **e**. Top, schematics of *Setdb1* genomic regions after degron engineering, with genotyping primers used for screening engineered clones indicated. Bottom, Genotyping of successfully targeted clones using primer pairs indicated at the top. The expected sizes are detailed in the supplementary table and was denoted with a red arrow. **f**. western blot of Setdb1 and loading control, αTubulin, of parental cTKO ESCs, S1SD and degron engineered dTKO mouse ESCs. The molecular weight in kilodalton (KD) is indicated on the left, and the expected band of wildtype Setdb1 and dTAG-Setdb1 fusion proteins are indicated with red arrows on the right. S1SD, mouse ESC line constitutively expressing 3HA-dTAG-Setdb1 fusion protein.

**Supplementary Fig. 3:**
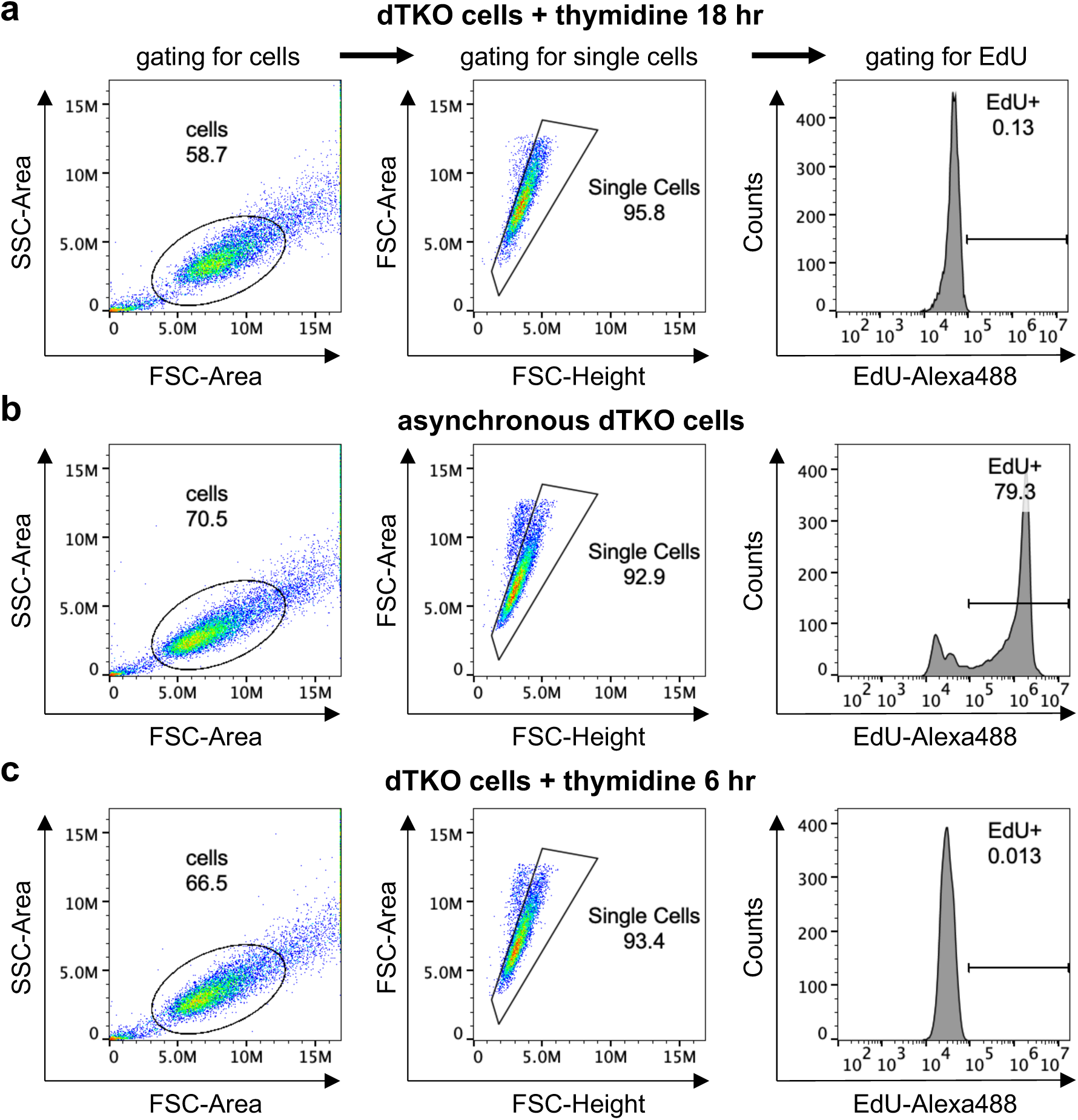
Gating strategies for FACS data. **a-c**, FACS analysis of EdU integration in mouse ESCs treated with thymidine for 18 hr (**a**), asynchronized dTKO (**b**), and dTKO cells treated with thymidine for 18 hr 6 hr (**c**). Cells were first gated based on FSC and SSC scatter plot to filter out cell debris (left), followed by gating for single cells that shows expected linear correlations between FSC-Height and FSC-Area (middle). The EdU-Alexa 488 fluorescent signal in single cell population was then analyzed (right). DNA synthesis in cells treated with 18 hr thymidine were blocked (**a**) were therefore were used as negative control to gate for EdU positive cells in **b,c**.

**Supplementary Fig. 4:**
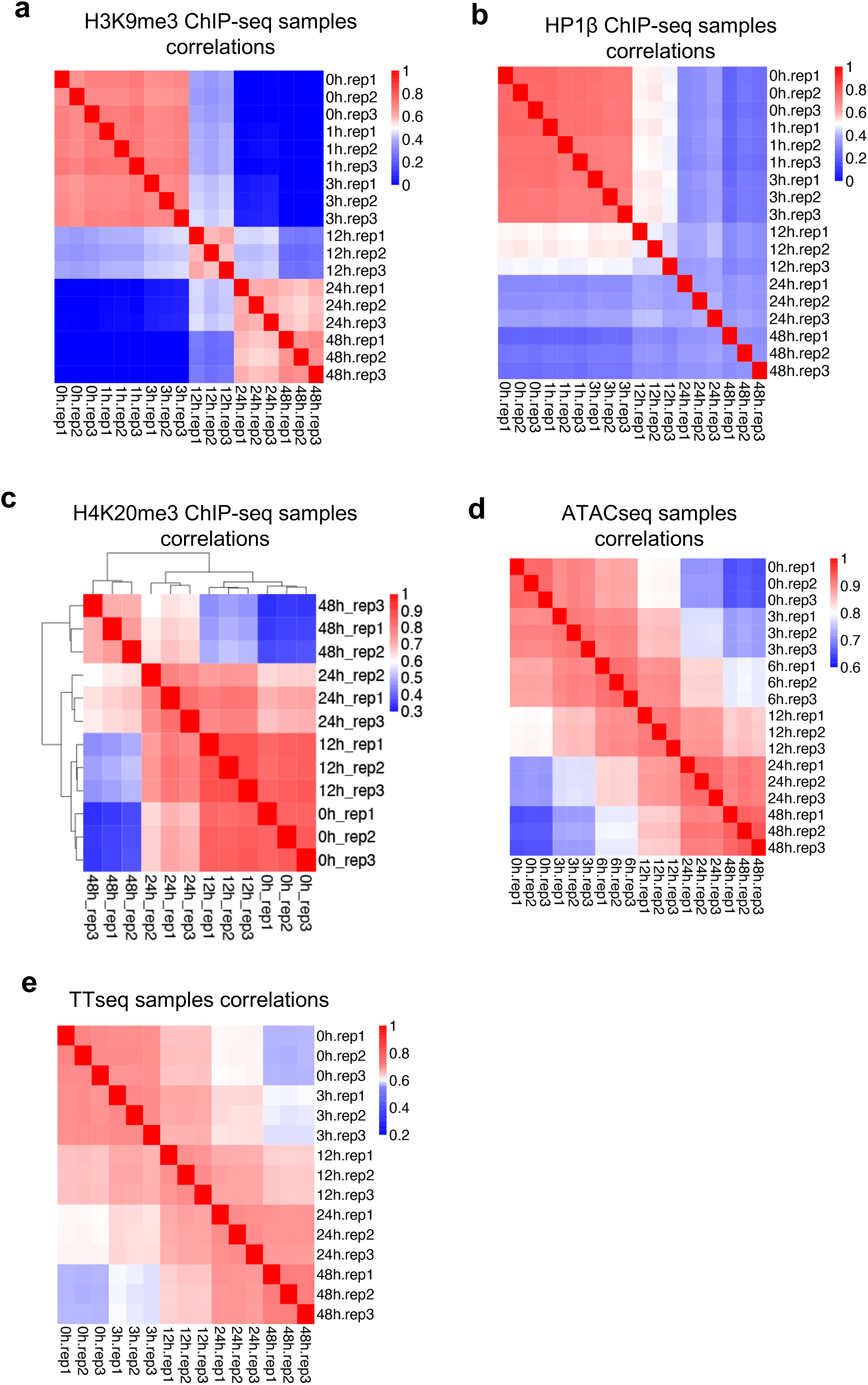
Reproducibility of ChIP-seq, ATAC-seq and TT-seq series. **a-e,** Heatmaps showing the spearman correlations between different samples in H3K9me3 (**a**), HP1β (**b**) and H4K20me3 (**c**) ChIP-seq series, ATAC-seq (**d**), and TT-seq series (**e**).

**Supplementary Fig. 5:**
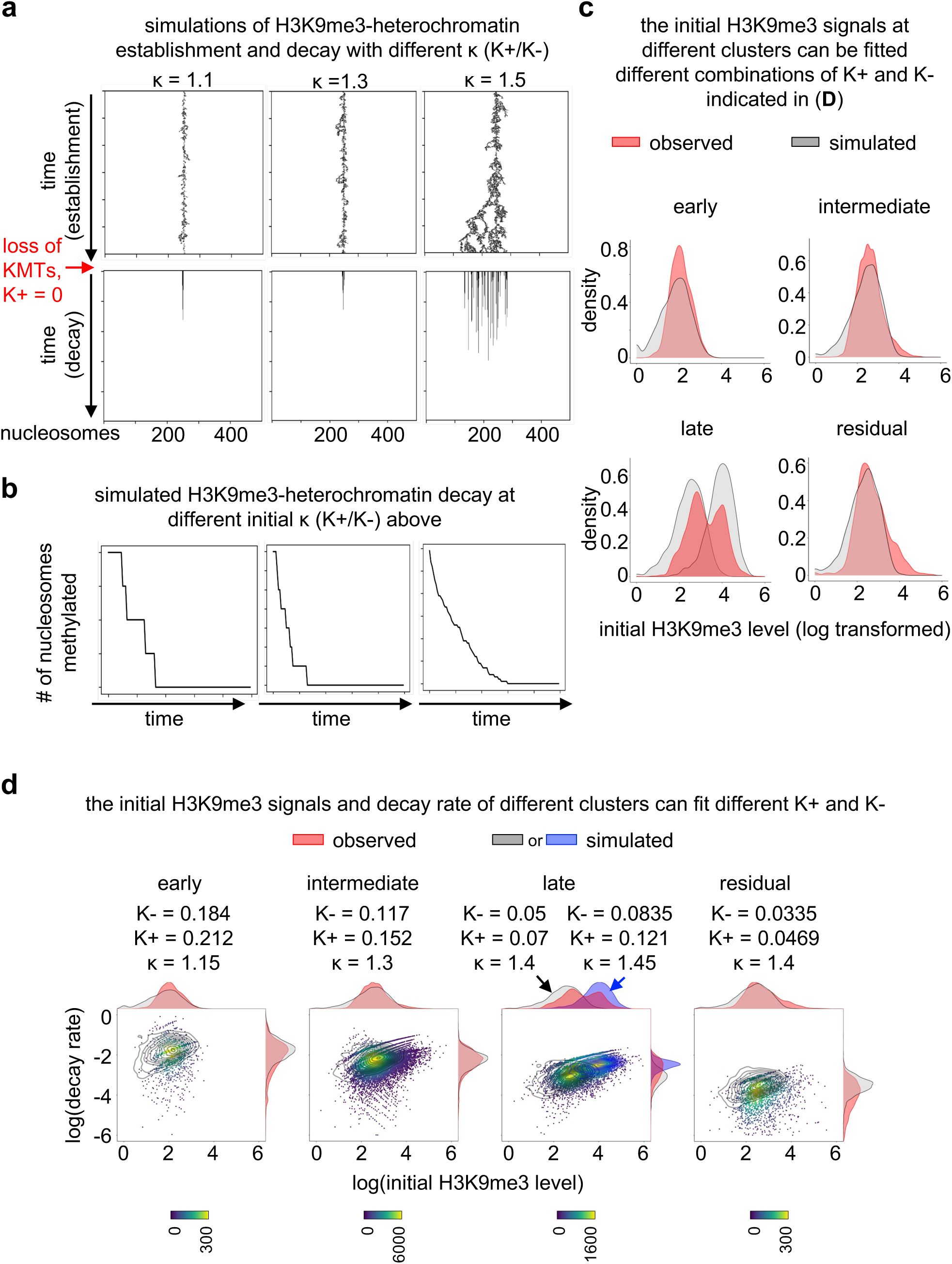
Mathematical modeling of H3K9me3 establishment and decay. **a**. Representative images of H3K9me3 traces in the simulated H3K9me3 establishment and decay at various κ (K+/K-). **b**. Representative images of the quantifications of H3K9me3 levels during the simulated H3K9me3 decay in the panel **a** above. **c**. Density plots showing the distribution of observed H3K9me3 signals (red) at different H3K9me3 clusters and the simulated H3K9me3 signals (grey) produced with the best fitted K+ and K- (and κ) parameters for each cluster. **d**. Dot plots showing the distribution of observed initial H3K9me3 signals and decay rate at different H3K9me3 clusters, super-imposed with a contour plot (in grey) showing the distributions of H3K9me3 signals and decay rate produced by the simulation with the model K+ and K- parameters indicated above. The density plot at the top and on the right shows the density plots showing the distributions of the initial H3K9me3 levels (top) and H3K9me3 decay rate (right) in the observed (red) and simulated data (grey or blue). Note that there are two sub-clusters in the late-H3K9me3 cluster, fitted with different K- parameters.

**Extended Data 1:**
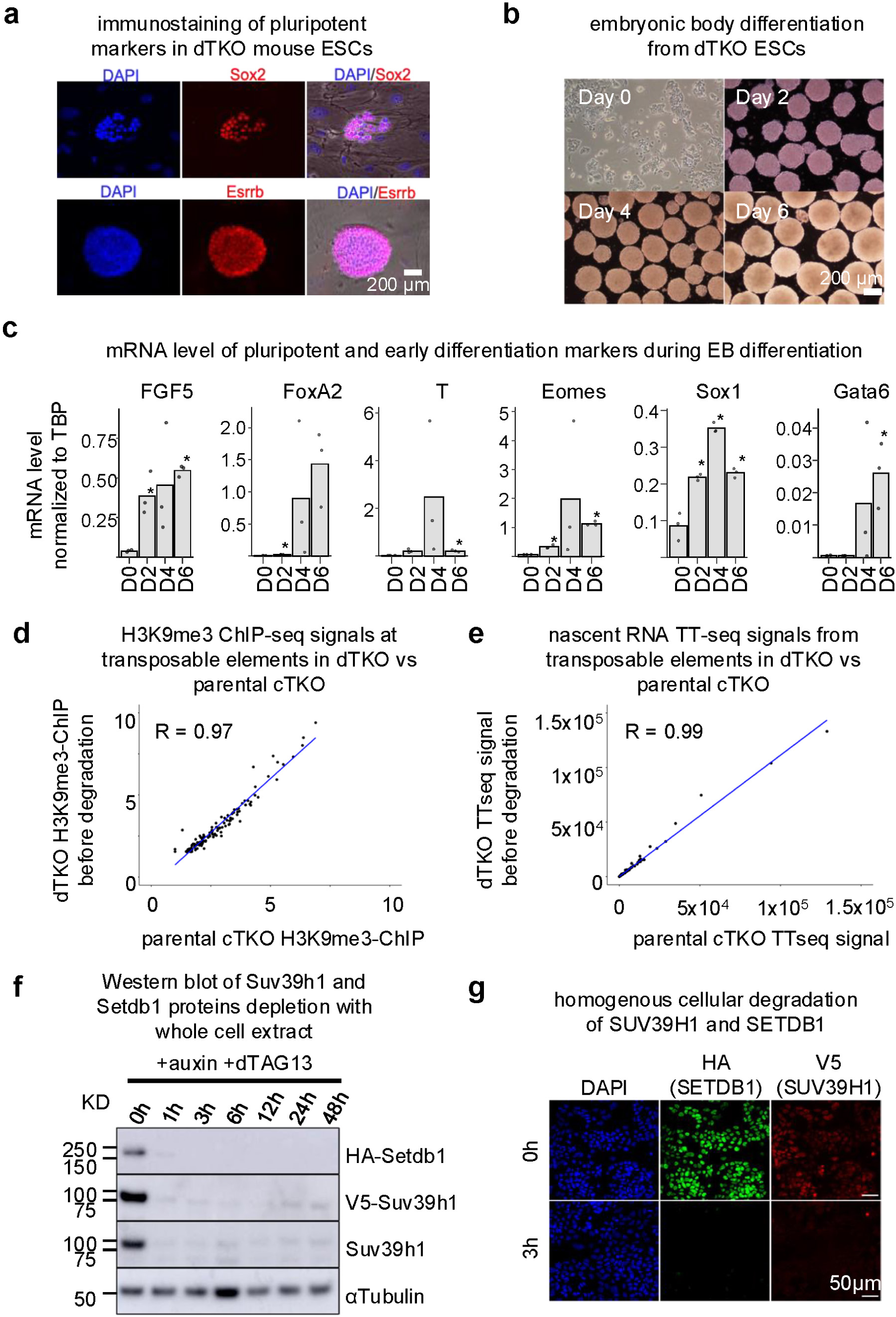

**Extended Data 2:**
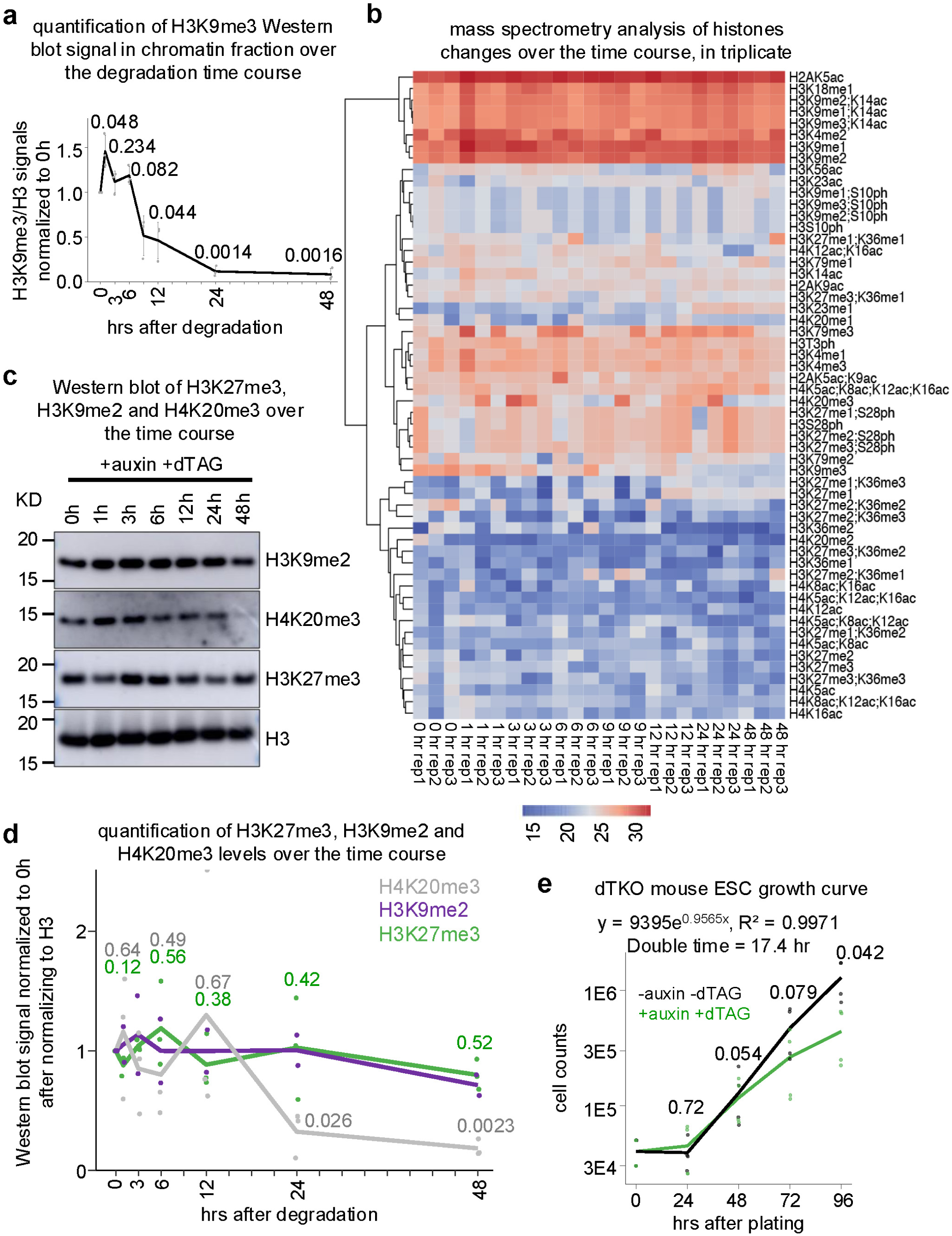

**Extended Data 3:**
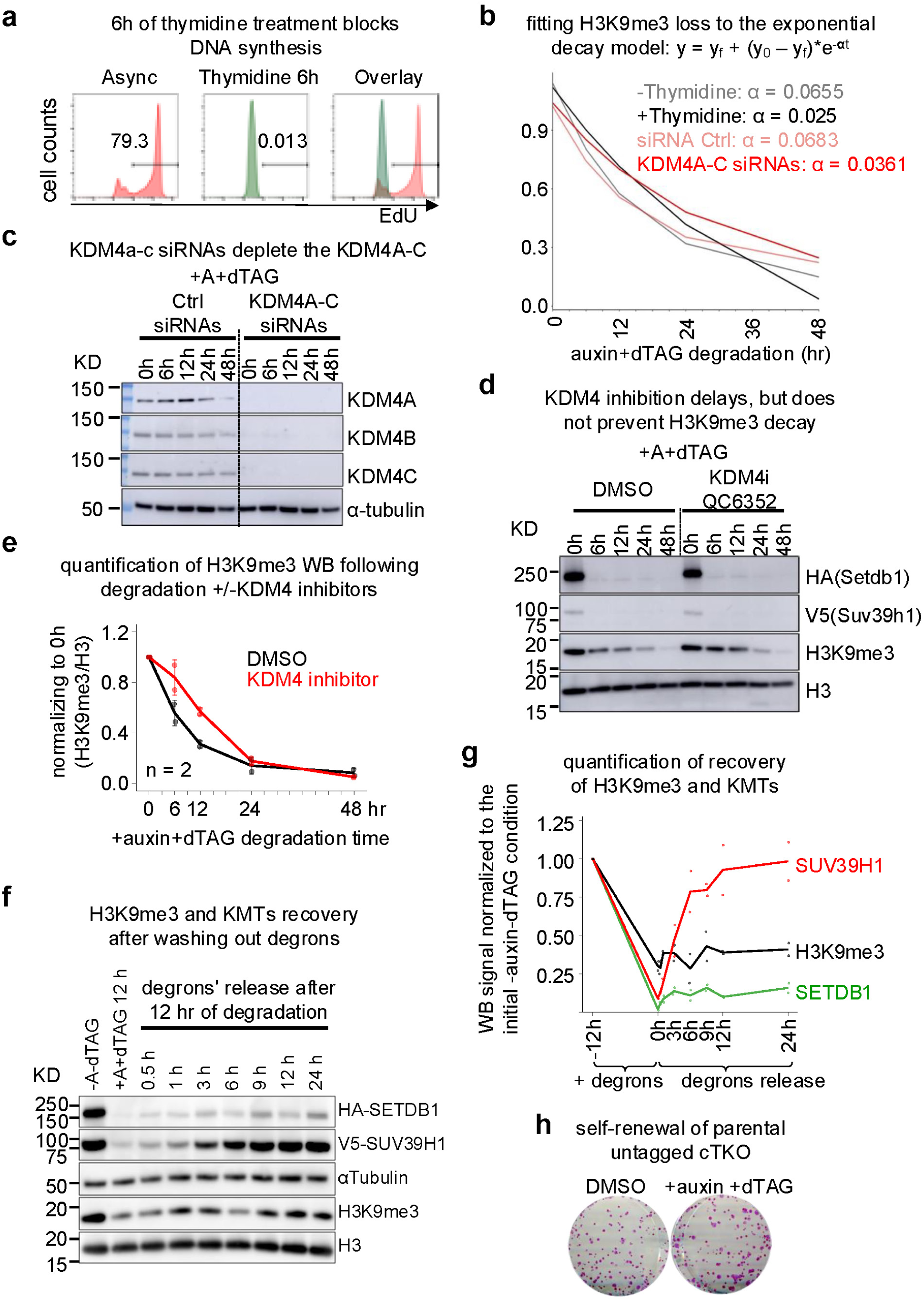

**Extended Data 4:**
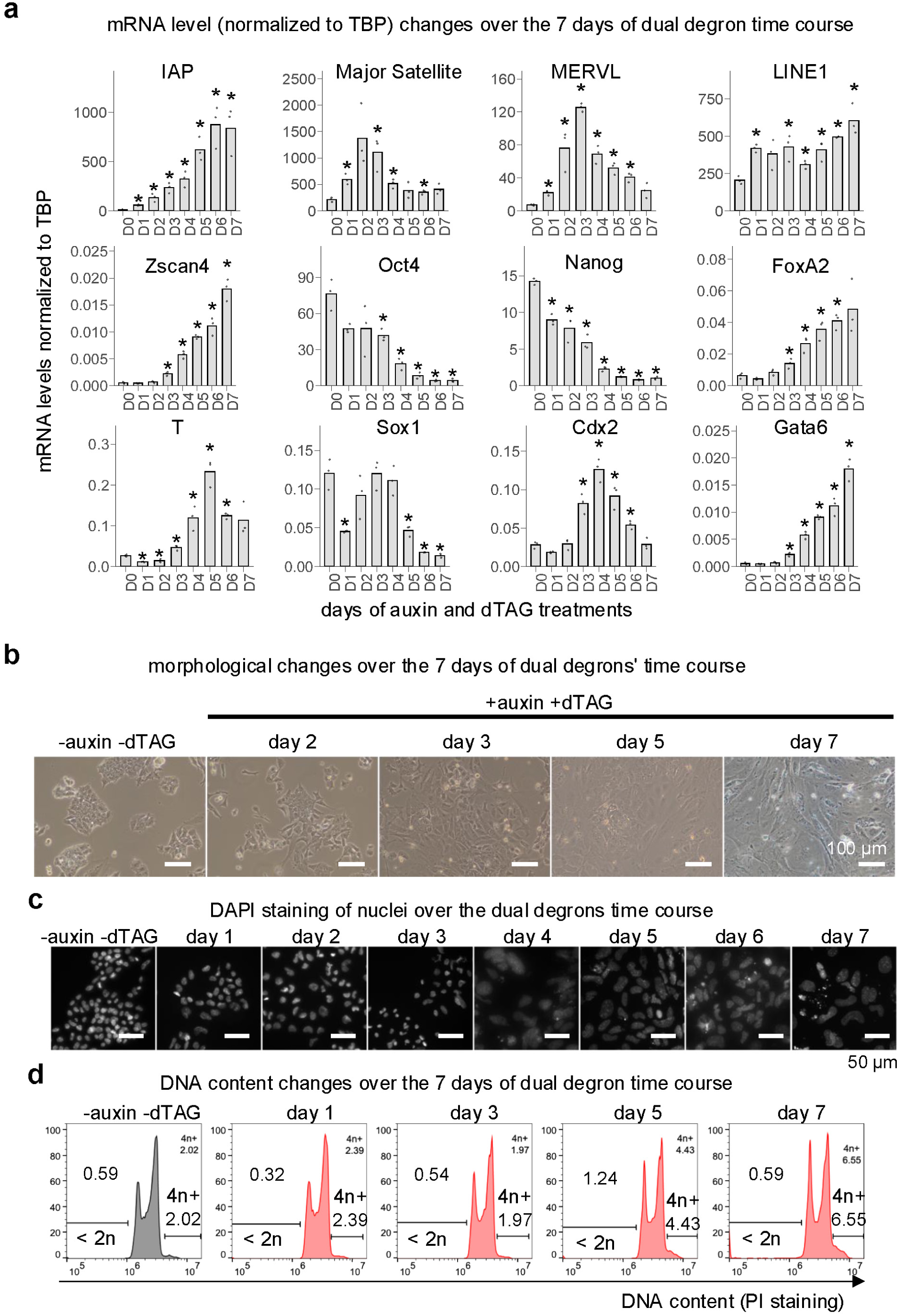

**Extended Data 5:**
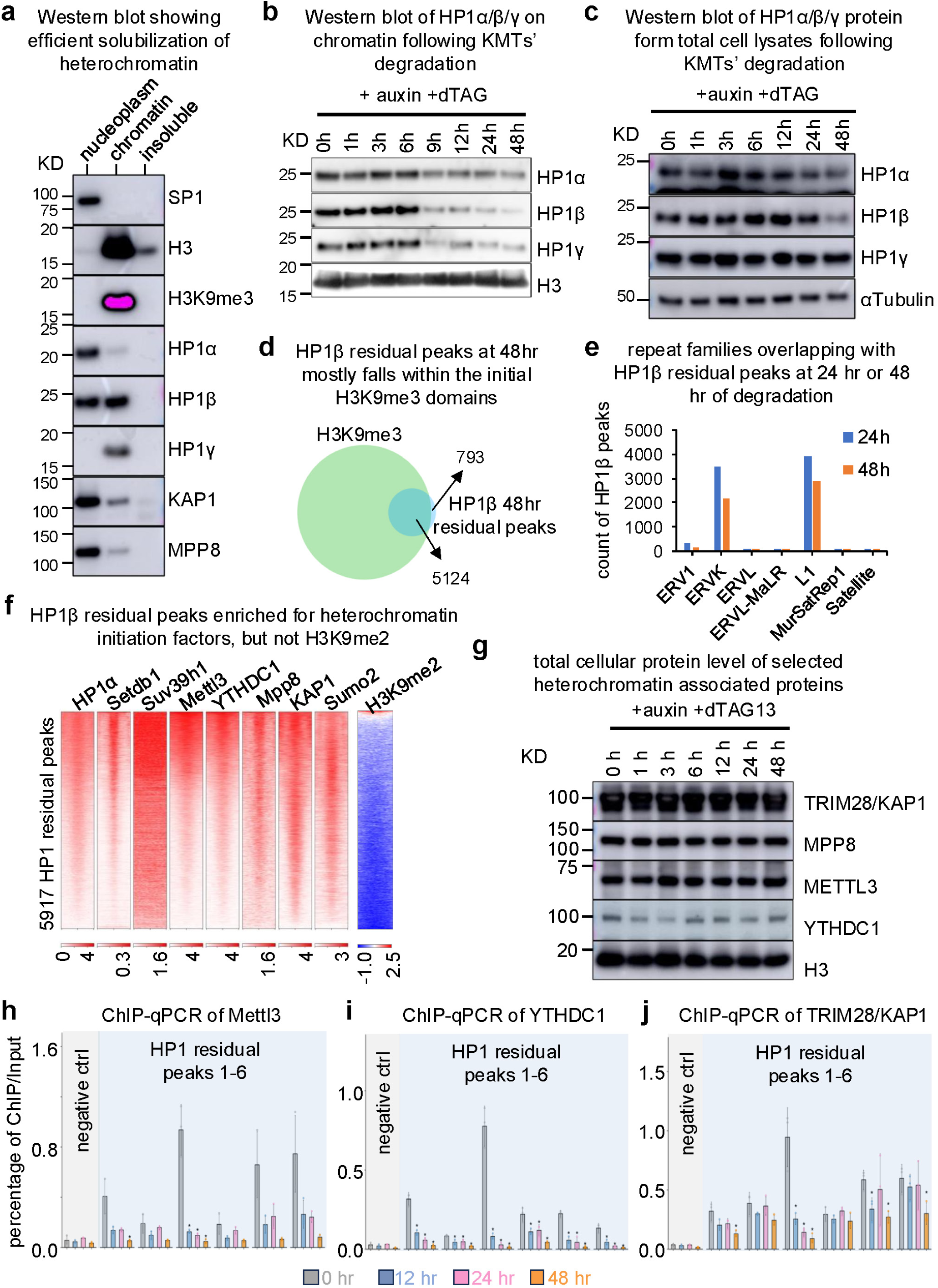

**Extended Data 6:**
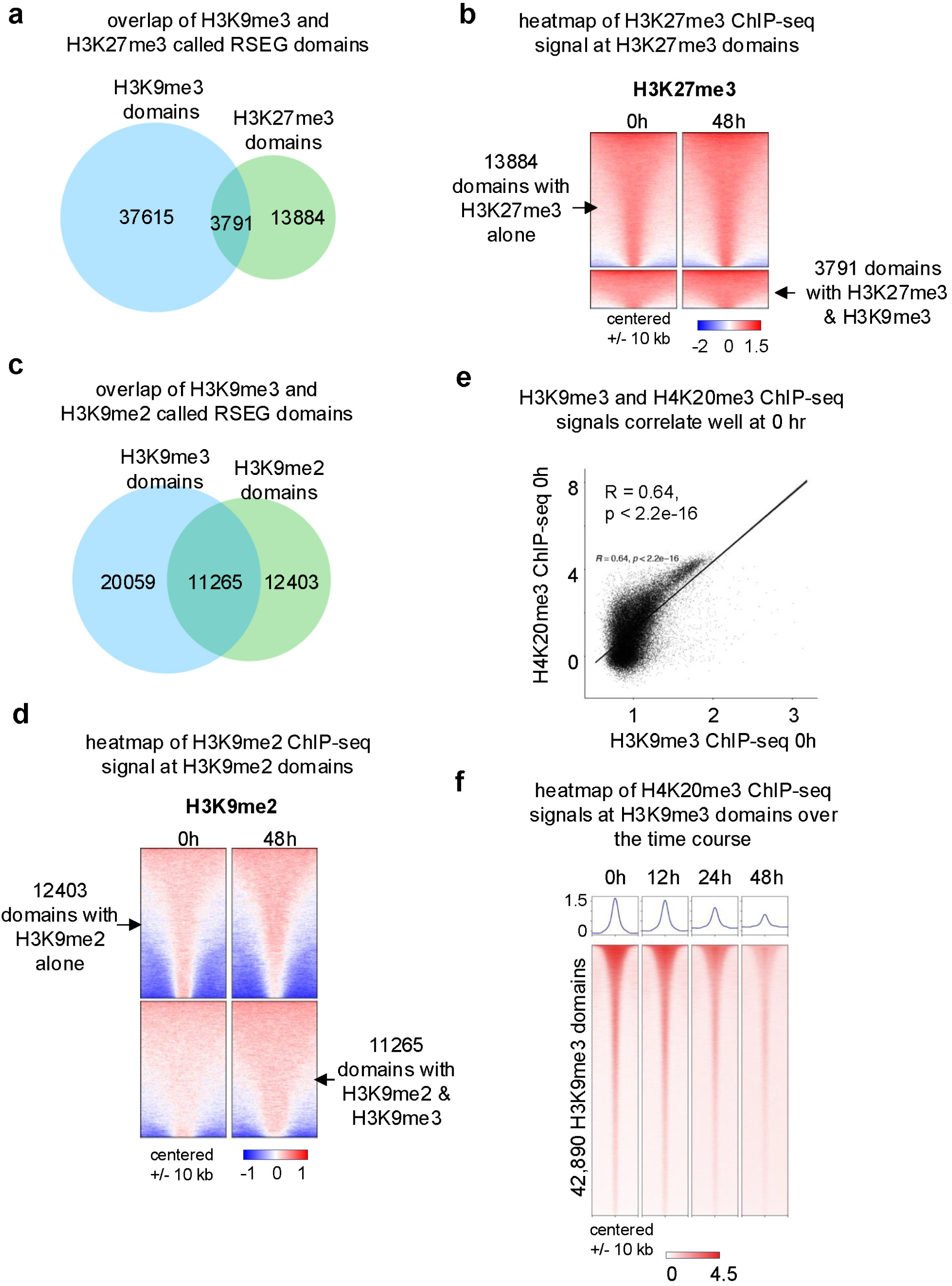

**Extended Data 7:**
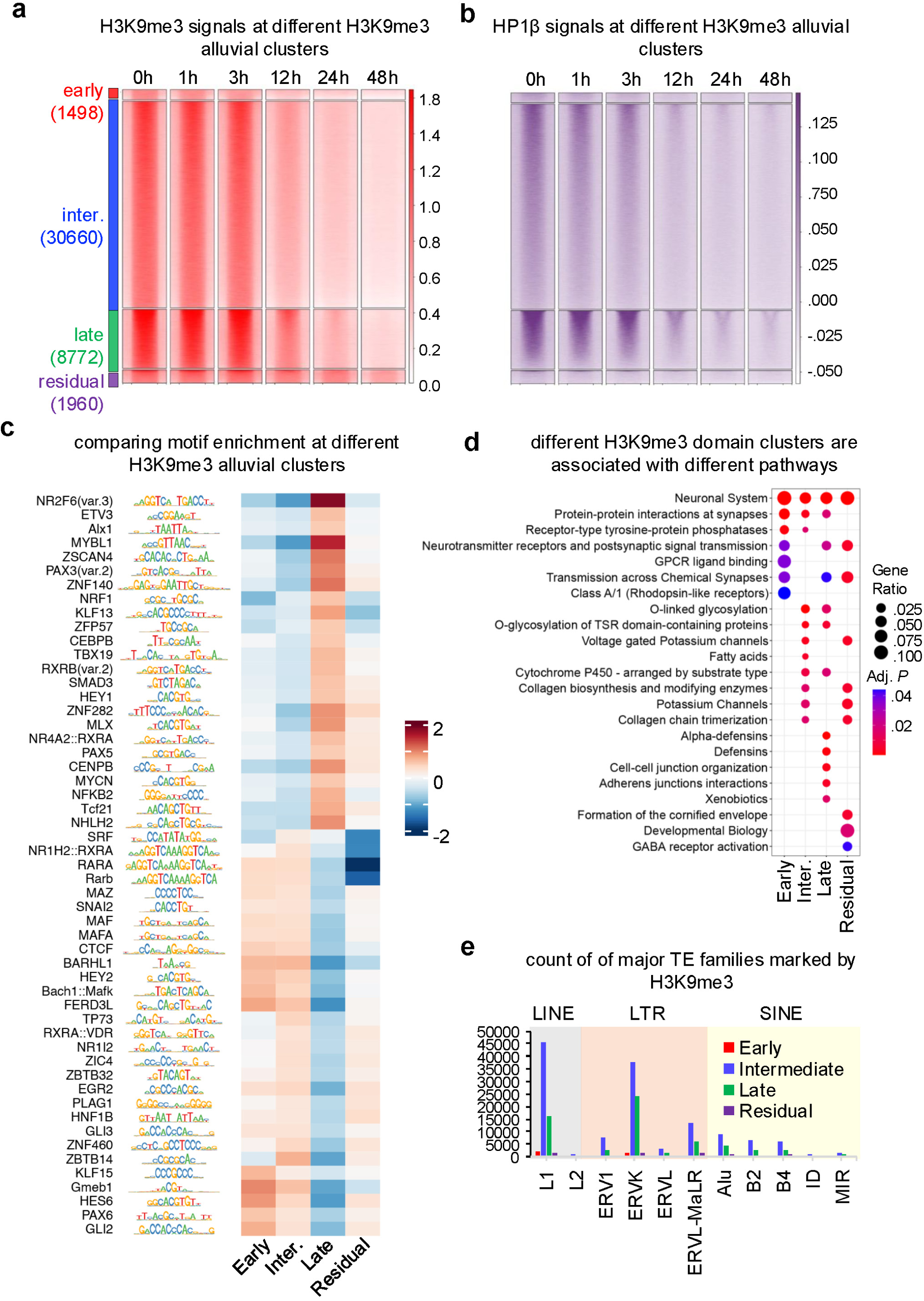

**Extended Data 8:**
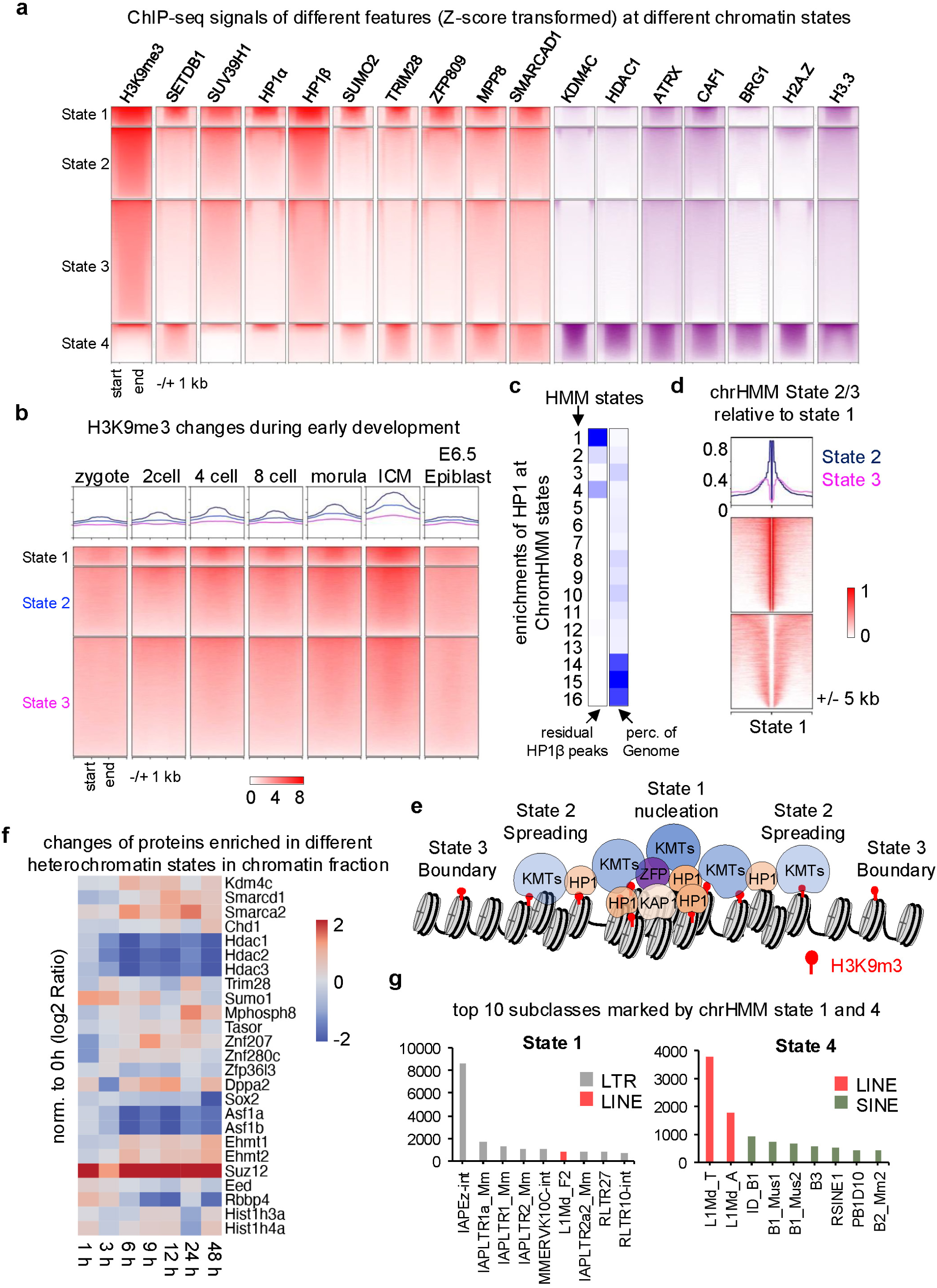

**Extended Data 9:**
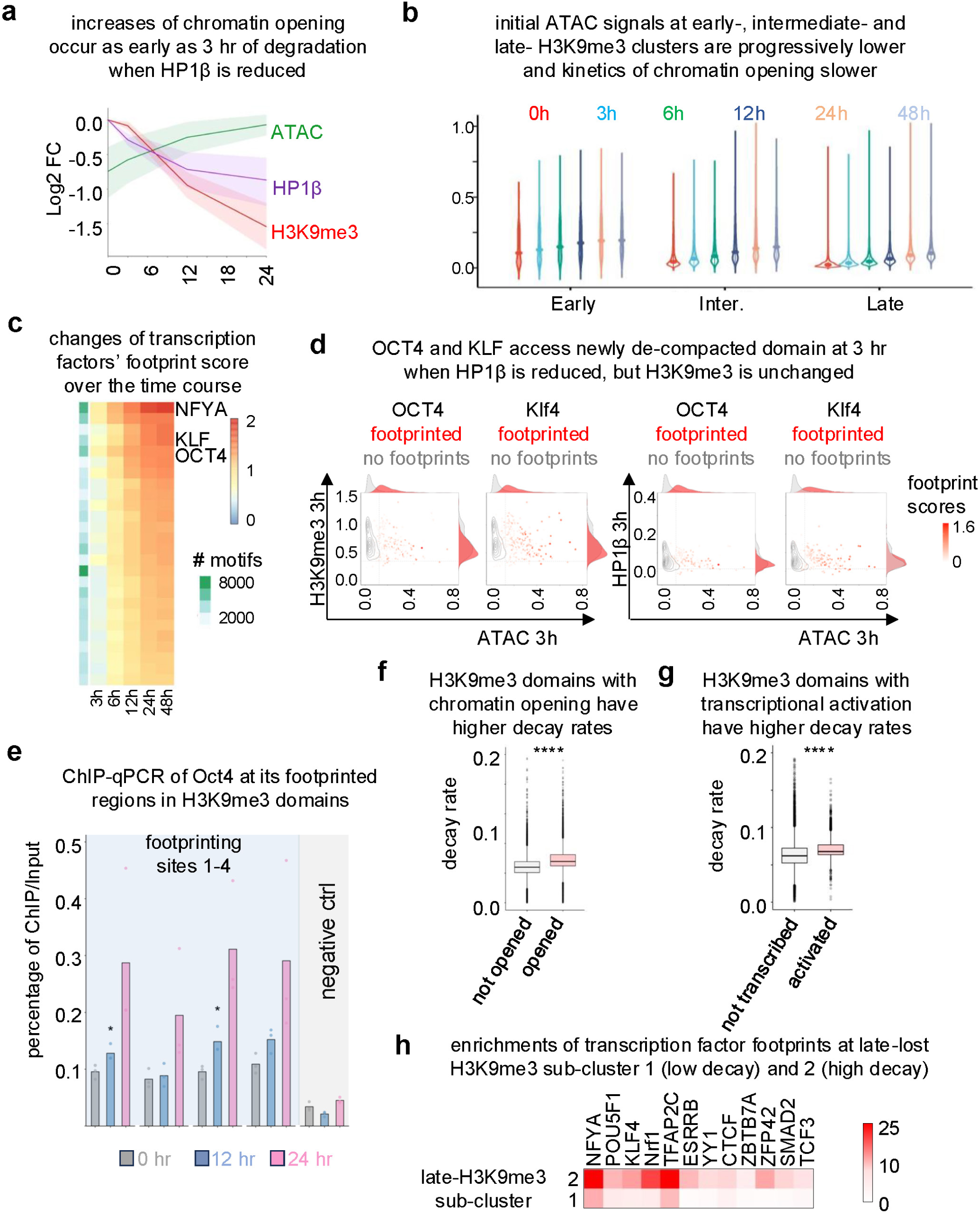

**Extended Data 10:**
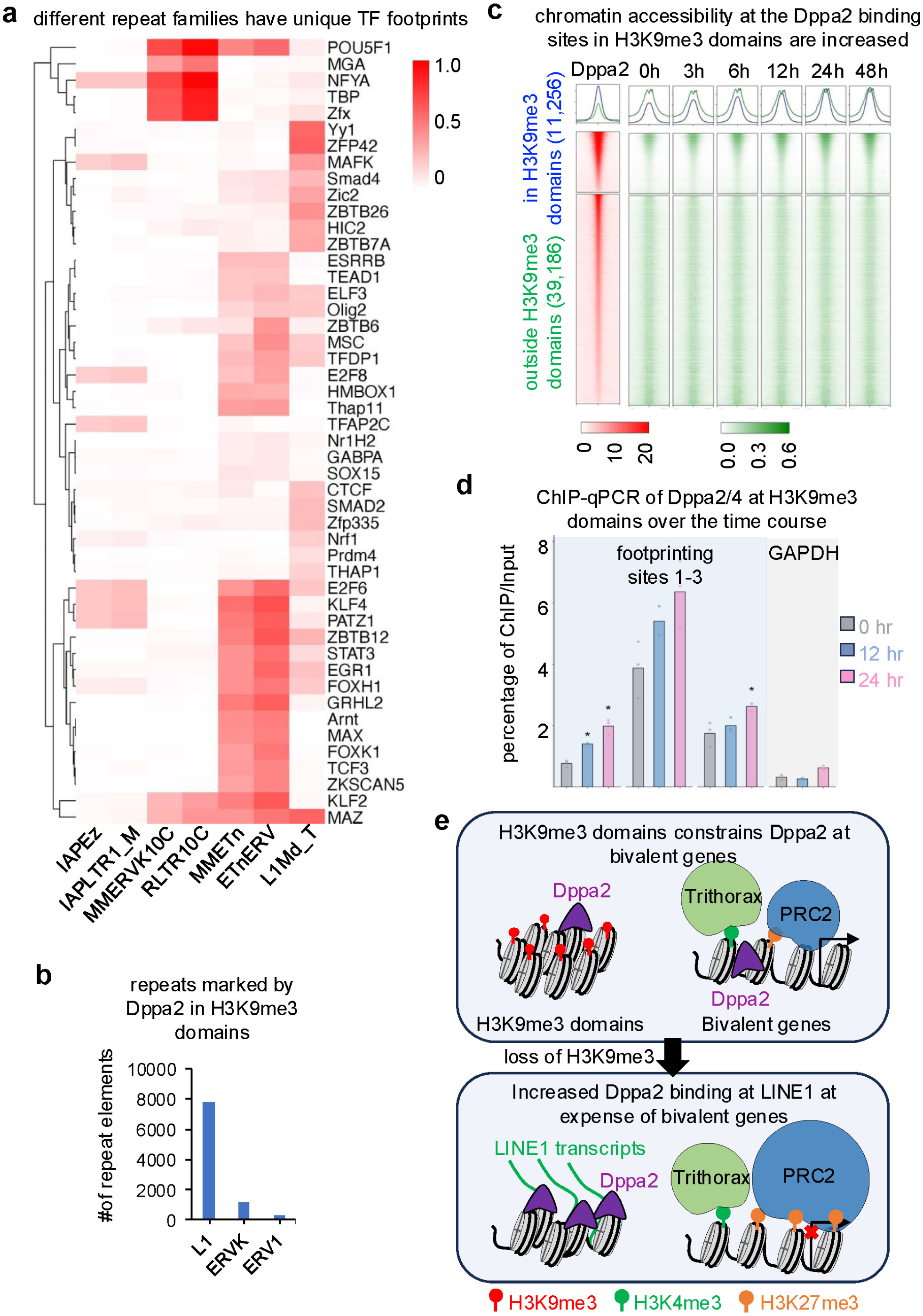

